# Dark matter of an orchid: metagenome of the microbiome associated with the rhizosphere of *Dactylorhiza traunsteineri*

**DOI:** 10.1101/2025.07.21.665863

**Authors:** Gabriel A. Vignolle, Leopold Zehetner, Christian Zimmerman, Domenico F. Savio, Ovidiu Paun, Robert L. Mach, Astrid R. Mach-Aigner, Julien Charest

## Abstract

Plant microbiota forms complex co-associations with its host, promoting health in natural environments. Root-associated bacteria (RAB) colonize root compartments and can modulate plant functions by producing bioactive compounds as part of their secondary metabolism. We present an in-depth analysis of the rhizosphere-associated microbiome of the endangered marsh orchid *Dactylorhiza traunsteineri*. Using deep sequencing of 16S rRNA genes, we identified *Proteobacteria*, *Actinobacteria*, *Myxococcota*, *Bacteroidota*, and *Acidobacteria* as predominant phyla associated with the rhizosphere of *D. traunsteineri*. Using deep shotgun metagenomics and de novo assembly, we extracted high-quality metagenome-assembled genomes (MAGs), revealing significant metabolic and biosynthetic potentials of the *D. traunsteineri*’s RABs. Our study offers a comprehensive investigation into the microbial community of the *D. traunsteineri* rhizosphere, highlighting a potential novel source of critical bioactive substances. Our study offers novel insights and a robust platform for future investigations into *D. traunsteineri*’s rhizosphere, crucial for understanding plant-microbe interactions and aiding conservation efforts for endangered orchids.

## INTRODUCTION

Every large multicellular organism studied so far, including plants, interacts with a plethora of microorganisms in its environment. The plant microbiota, which includes bacteria, fungi, protists, nematodes, and viruses, can form complex co-associations with its host. Plant microbial communities have been shown to promote plant growth, nutrient uptake, as well as pathogen resistance, thus promoting plant health in its natural environment (Richardson and Simpson, 2011, Pieterse et al., 2014, Trivedi et al., 2016, Backer et al., 2018, Gouda et al., 2018, Trivedi et al., 2020). However, our understanding of plant-microbial alliances’ functional mechanisms and broad ecological implications is still in its early stages (Martin et al., 2017).

Among the bacterial communities interacting with plants, a subset known as the root-associated bacteria (RAB) colonizes root compartments, such as the rhizosphere (i.e. the narrow zone surrounding and influenced by plant roots growth and metabolic activity) and intraradical regions and has the ability to modulate plant functions (Mendes et al., 2013, Qu et al., 2020, Dobbelaere et al., 2003, Walker et al., 2003). It has been suggested that the holobiont, defined as the host and its associated microbiome, forms a discrete symbiotic ecological unit that has co-evolved to maintain host function and fitness (Hassani et al., 2018, Uroz et al., 2019, Vandenkoornhuyse et al., 2015, Trivedi et al., 2020, Liu et al., 2019). Plants selectively recruit RAB depending on different factors such as host genotype, metabolic profiles, root exudates, and soil physicochemical properties (Bulgarelli et al., 2013, Hassani et al., 2018, Liu et al., 2019, Uroz et al., 2019). Although the plant and its exudates mostly influence this microenvironment, environmental influences also significantly shape its composition (Vives-Peris et al., 2020, Najera et al., 2020). Furthermore, diverse endophytic bacteria possess the ability to produce bioactive compounds of usually low molecular mass, with a wide range of reported functions, as part of their secondary metabolism. These compounds, which are not crucial for the fundamental cellular processes of bacteria in controlled laboratory conditions, often confer selective advantages within natural ecosystems. Such secondary metabolites (SMs) include antibiotics, pigments, toxins, enzyme inhibitors, immunomodulators, effectors of ecological competition or symbiosis, as well as compounds with hormonal activity or particular effects on lipids or carbohydrate metabolism (reviewed in (Fouillaud and Dufosse, 2022)). Many RABs have been reported to enhance plant growth through processes such as nutrient acquisition, hormonal regulation, protection against pathogens, and mitigation of abiotic stresses (reviewed and discussed in (Narayanan and Glick, 2022, Glick, 1995, Olanrewaju et al., 2017, Glick, 2012, Hassani et al., 2018, Mendes et al., 2013, Qu et al., 2020, Trivedi et al., 2020, Uroz et al., 2019, Vandenkoornhuyse et al., 2015)). These RAB, conferring growth benefits to plant hosts, are collectively referred to as plant growth-promoting rhizobacteria (PGPR), which typically belong to the *Proteobacteria*, *Actinobacteria*, *Firmicutes*, and *Bacteroidetes* phyla (Santoyo et al., 2016). Within these phyla, genera such as *Rhizobium*, *Bacillus*, *Pseudomonas*, and *Burkholderia* are prominently recognized as PGPR and have demonstrated functional contributions in mediating seed germination, plant growth and biocontrol of plant disease, notably by conveying secondary metabolites inclusive of antibiotics, volatile compounds, phytohormones and siderophores (Glick, 2012, Kang et al., 2012, Walia et al., 2013, Bloemberg and Lugtenberg, 2001). Notably, the publication of whole-genome sequences of rhizosphere-inhabiting bacteria facilitates the identification of genes involved in the regulation and production of SMs by comparative and functional genomics. However, despite the established knowledge that symbiotic associations between roots and PGPR provide multiple growth benefits to host plants, little is known regarding the role of PGPR and other RAB in shaping plant ecology through their secondary metabolism (Siefert et al., 2019, Kaur and Sharma, 2021).

Orchids, which account for approximately 10% of all seed plants, exhibit intricate associations with diverse microorganisms in their natural habitat (Givnish et al., 2016, Givnish et al., 2015). One example is mycoheterotrophy, an alternative nutritional strategy observed in certain plant species, particularly orchids, which derive sugars and essential nutrients from symbiotic associations with mycorrhizal fungi (Timilsena et al., 2023). The marsh orchid *Dactylorhiza traunsteineri* is an allotetraploid (2*n* = 80), mainly found in the eastern hemisphere (Paun et al., 2010). *Dactylorhiza traunsteineri* is part of the *D. majalis* complex, which has evolved polytopically from unidirectional hybridization between the diploid species *Dactylorhiza fuchsii* and *D. incarnata* (Pillon et al., 2007, Brandrud et al., 2020). It is slender, usually low-grown with narrow leaves and a lax few-flowered inflorescence, has narrow soil moisture and pH tolerance, and grows in calcareous fens, typically on naturally open sites associated with seepage zones (Kull and Hutchings, 2006, Soó, 1980, Baumann and Künkele, 1988, Delforge, 2001). The species occurs in low-nutrient soils, characterized in particular by very low levels of available nitrate and other macro and micronutrients (Wolfe et al., 2023). A recent study documented, at two different localities, a high diversity of fungi associated with the roots of *D. traunsteineri* in its natural environment, in comparison to its sibling *D. majalis* (Emelianova et al., 2024). Although hybrids and allopolyploids are known to be more resistant to environmental changes due to their larger genomic potential, *D. traunsteineri* is classified as endangered by several organizations in Europe and has proven challenging to grow and maintain under artificial environmental conditions, which is a requirement for ex-situ conservation efforts (Paun et al., 2007, Blinova and Uotila, 2012, Bartók et al., 2018, Paun et al., 2011).

This study presents a comprehensive analysis of the rhizosphere-associated bacterial microbiome of *Dactylorhiza traunsteineri*. Our community analysis, based on 16S ribosomal DNA (rDNA)-targeted sequencing, identified *Proteobacteria*, *Actinobacteria*, *Myxococcota*, *Bacteroidota*, and *Acidobacteria* as the predominant phyla within the *D. traunsteineri* rhizosphere. Leveraging deep shotgun metagenomics and de novo metagenome assembly, we extracted 47 MAGs, unveiling significant metabolic and biosynthetic capabilities of the orchid’s RAB. Notably, this study represents the first comprehensive investigation into the microbial community of the *D. traunsteineri* rhizosphere, highlighting a potential novel source of critical bioactive substances. Our findings illuminate previously uncharacterized PGPR SM pathways and the discovery of “dark matter” biosynthetic gene clusters, underscoring the rhizosphere’s untapped potential for biotechnological and ecological applications. This research advances our understanding of *D. traunsteineri*’s microbial interactions and sets the stage for future studies to exploit these microbial consortia for conservation and bioprospecting efforts.

## MATERIALS & METHODS

### Sampling and sample processing

*Dactylorhiza traunsteineri* individuals (from the *turfosa* subspecies) were isolated from the Kitzbühel alpine town region, in the western Austrian province of Tyrol on the 25^th^ of May 2018. The average temperatures recorded in this region during summer are between 13°C and 26°C, and between 0°C and 6°C during winter, with an annual total rainfall estimated to 1,443 mm, the majority occurring primarily during summer in June (average 188 mm). The individuals were potted and maintained at the Botanical Garden of the University of Vienna under conditions mimicking those of the sampling region. The sampling of the rhizosphere soil was performed on 03.02.2020 under sterile laboratory conditions. The plantlets were carefully uprooted, the soil discarded by shaking the roots, and the adherent soil directly in contact with the root system collected. Additionally, the roots of the plantlets were washed for 5 min with sterile DNA-free laboratory-grade Sartorius avium® filtered water. Samples were collected in sterile 50 ml Cellstar tubes. Both samples were shock-frozen in liquid nitrogen and kept at -80°C until the metagenomic DNA extraction was performed.

### Metagenomic DNA extraction

Isolation of the metagenomic DNA was performed from the rhizosphere soil and the plantlet wash lyophilized samples in a sterile environment. The root wash samples were lyophilized using the FreeZone Freeze Dryer system [LabConco; Cat. 700201000]. 500 mg of lyophilized mass from each sample was transferred to a 2 mL screw cap reaction tube with sterile glass beads. Samples were first disrupted using the FastPrep-24 Classic homogenizer [MPBio; Cat. 116004500] at setting level 6 for 30 seconds. 1 mL of 2xCTAB buffer (pH 8.0) [100 mM Tris (pH 8.0); 20 mM EDTA (pH 8.0); 1.4 M NaCl; 2% (w/v) CTAB] with 4 mL β-mercaptoethanol, was preheated at 55°C and added to the samples, before disrupting the biomass a second time using the FastPrep-24 Classic homogenizer, twice for 30 seconds at setting level 5. The samples were then incubated for 20 minutes at 65°C and centrifuged for 1 minute at 10,000 rpm before transferring the supernatant and foam to fresh, sterilized 2 mL reaction tubes. Next, 400 µL Phenol and 400 µL SEVAG [4% (v/v) isoamyl alcohol; 96% (v/v) chloroform] were added to the samples before vigorous mixing and incubation for 1 hour at room temperature. The aqueous phases were transferred to fresh, sterilized 2 mL reaction tubes, and 800 µL chloroform was added to the samples. The samples were mixed vigorously and centrifuged for 1 minute at 10 000 rpm, before transferring the aqueous phase to fresh sterilized 2 mL reaction tubes. 2 µL RNase A [10 mg/ml [ThermoFisher; Cat: EN0531]] were then added, and the samples mixed by inversion and incubated for 15 minutes at 60°C.

Next, 2 volumes of ice-cold 96% ethanol were added to the samples and incubated overnight at -20°C for RNase A inactivation and DNA precipitation. Following overnight incubation, the samples were centrifuged for 20 minutes at 10,000 rpm at 4°C. The supernatants were discarded, and the DNA pellets were washed once in 500 µL 96% ethanol. The samples were then centrifuged for 10 minutes at 10,000 rpm at 4°C, and the DNA pellets washed twice with 500 µL 70% ethanol. After centrifuging for an additional 10 minutes at 10,000 rpm at 4°C, the DNA pellets were dried and resuspended in 100 µL ultra-pure water and incubated overnight at 4°C, before quality assessment. Samples were stored at 4°C.

### 16*S* rDNA amplicon sequencing and analysis

For bacterial community composition analysis, the V3-V4 fragment of the 16S rDNA gene was amplified from the genomic DNA extracted from the soil and root wash samples. 16S rRNA fragment concentrations in the DNA extracts were quantified using domain-specific quantitative PCR (Fontaine et al., 2023). DNA extracts were normalized based on the 16S rRNA gene concentrations to ensure 16S rRNA gene template molarity for amplification and a two-step barcoding procedure. A first amplification reaction was performed using the 16S rDNA 341F and 805R primer set, containing adapters for the introduction of Illumina adapters (341F + adapter: ACACTCTTTCCCTACACGACGCTCTTCCGATCTNNNNCCTACGGGNGGCWGCAG; 805R + adapter: AGACGTGTGCTCTTCCGATCTGACTACHVGGGTATCTAATCC), using Q5 high-fidelity DNA polymerase [New England Biolabs; Cat: M0491] and the following cycling conditions: 98°C for 1 minute, followed by 20 cycles of 98°C for 10 seconds, 62°C for 30 seconds, 72°C for 30 seconds, and a final extension at 72°C for 2 minutes. The 16S rRNA gene copy concentrations in DNA extracts were determined through quantitative PCR and normalized to equal 16S rRNA fragment copy number to enhance comparability and reduce PCR bias, and purified using the Agencourt AMPure XP [Beckman; Cat: B23319] purification system. Sequencing indexes were introduced in a second amplification reaction with sample-specific barcodes (FWD: AATGATACGGCGACCACCGAGATCTACAC-[index]-ACACTCTTTCCCTACACGACG; REV: CAAGCAGAAGACGGCATACGAGAT-[index]-GTGACTGGAGTTCAGACGTGTGCTCTTCCGATCT) using Q5 high-fidelity DNA polymerase and the following cycling conditions: 98°C for 1 minute, followed by 15 cycles of 98°C for 10 seconds, 66°C for 30 seconds, 72°C for 30 seconds, and a final extension at 72°C for 2 minutes. Amplicon libraries were purified using the Agencourt AMPure XP purification system and quantified using PicoGreen kit [ThermoFisher; Cat: P7589]. The libraries were pooled and diluted to 4 nM before being sequenced for 300 cycles pair-end on the Illumina MiSeq system using the MiSeq® Reagent Kit v2 [Illumina; Cat: MS-102-2002].

Following sequencing, the quality of the 16S amplicon sequences was verified using FastQC (v.0.11.9) (Andrews, 2010) and sequencing adapter sequences, as well as low-quality reads, were removed from the raw read data using Trimmomatic v.0.40 (Bolger et al., 2014) with the following parameters: *ILLUMINACLIP:TruSeq3-PE-2.fa:2:30:10:2:keepBothReads LEADING:3 TRAILING:3 SLIDINGWINDOW:4:15 MINLEN:36*. 16S amplicon sequencing data analysis has been performed using the R package dada2 (v.1.26.0) (Callahan et al., 2016), which includes quality filtering, dereplication, dataset-specific error model determination, ASV interference, chimeral removal, as well as taxonomic assignment, as part of its standard workflow.

### Shotgun Sequencing

Shotgun metagenomics sequencing was conducted on the genomic DNA extracted from the soil and root samples. Evaluation of the initial DNA concentrations in the prepared samples was carried out using the Qubit dsDNA HS Assay Kit [ThermoFisher; Cat: Q32851]. The libraries were prepared using 1000 ng of genomic DNA, which was fragmented using the Bioruptor sonication device [Diagenode; Cat: B01020001]. Fragments were cleaned up using the GeneJet PCR Purification Kit [ThermoFisher; Cat. #K0701] and size-selected for a 300 bp using Agencourt AMPure XP beads. Sequencing libraries were prepared following the standard NEBNext Ultra II DNA Library Prep with Sample Purification Beads [New England Biolabs; Cat: E7103] procedure. The final library concentrations were determined using the Qubit dsDNA HS Assay Kit and the average size of the library was determined using the Agilent 5200 Fragment Analyzer System [Agilent; Cat: M5310AA]. The libraries were pooled and diluted to 4 nM before being sequenced for 600 cycles pair-end on the Illumina MiSeq system using the MiSeq® Reagent Kit v3 [Illumina; Cat: MS-102-3303].

### Metagenome assembly, genome binning and quality assessment

The quality of the metagenomic reads was verified using FastQC (v.0.11.9) (Andrews, 2010) and sequencing adapter sequences, as well as low-quality reads, were removed from the sequenced data using Trimmomatic v.0.40 (Bolger et al., 2014) with the following parameters: *ILLUMINACLIP:TruSeq3-PE-2.fa:2:30:10:2:keepBothReads LEADING:3 TRAILING:3 SLIDINGWINDOW:4:15 MINLEN:36*. The quality trimmed reads from both samples were combined into one large dataset, containing the reads from both samples. The combined dataset was assembled using MEGAHIT (v.1.2.9) (Li et al., 2015) (parameters *--k-min 27 --k-max 247 --k-step 10*). Next, the paired-end metagenomic trimmed reads were mapped back to the generated contigs using bowtie2 (v.2.3.5.1) (Langmead and Salzberg, 2012) to generate coverage information, and sorted using SAMtools (v.1.10). To generate MAGs, contigs were binned using Autometa (v2.0) (Miller et al., 2019). Unclustered contigs were succinctly binned using MetaBAT2 (v.17) (Kang et al., 2015). For further improvement, the corresponding reads from each MAGs’ contigs were extracted and reassembled using SPAdes (v.3.14.1) (Nurk et al., 2017, Bankevich et al., 2012) using the meta option. Finally, the quality of each MAG was evaluated using CheckM (v.1.1.3) (Parks et al., 2015) and QUAST (v.5.0.2) (Mikheenko et al., 2018), to determine their completeness, contamination and heterogeneity.

### MAG analysis and taxonomical classification

For each MAG, ribosomal sequences were extracted based on the KEGG (Kanehisa, 2019, Kanehisa et al., 2023, Kanehisa and Goto, 2000) and PANNZER2 (Toronen et al., 2018) annotations, and quantified by type to identify the most abundant representatives in each MAG. For each binned MAG, taxonomic and lowest common ancestor predictions were generated using Autometa (Miller et al., 2019), based on the non-redundant NCBI database (O’Leary et al., 2016). Furthermore, a library containing genomic sequences of 199 reference organisms (Supp. Table 1), obtained from the NCBI database, was generated to calculate the average nucleotide identify (ANI) of each MAG (Supp. Figure 1) using FastANI (v.1.33) (Jain et al., 2018).

**Table 1.** Sequencing statistics (16S profiling)

| Samples | SRA accession no. | No. of raw<br>sequence reads | No. of reads that<br>passed quality<br>control | ASVs after<br>chimera removal |
| --- | --- | --- | --- | --- |
| Soil-R1 | SRR29434315 | 36,584 | 25,981 | 14,743 |
| Soil-R2 | SRR29434314 | 95,543 | 68,635 | 45,317 |
| Wash-R1 | SRR29434311 | 29,562 | 20,461 | 11,303 |
| Wash-R2 | SRR29434310 | 79,246 | 55,090 | 35,285 |

### Gene prediction and annotation

For each generated MAG, genes and proteins were predicted using Prodigal (v.2.6.3) (Hyatt et al., 2010), and the sequences annotated using the KEGG (Kanehisa, 2019, Kanehisa et al., 2023, Kanehisa and Goto, 2000) and PANNZER2 (Toronen et al., 2018) databases. The KEGG annotation was further used to investigate metabolic pathways, whereas PANNZER2 was used to annotate gene ontology and functional domains. Additionally, biosynthetic gene clusters (BGCs) for each MAG were predicted using antiSMASH (v.6.1.1) (Blin et al., 2017, Blin et al., 2021), DeepBGC (v.0.1.31) (Hannigan et al., 2019), and GECCO (v.0.9.10) (Carroll et al., 2021), annotated using the KEGG and PANNZER2 databases and evaluated for completeness.

## RESULTS

### 16S rRNA gene amplicon sequencing/taxonomic identification

To gain insights into the rhizosphere of *D. traunsteineri*, a potted orchid (Individuum 1 from Kitzbühel, sampled on 25.05.2018), maintained in the botanical garden of the University of Vienna (Universität Wien), has been transferred to our laboratory environment (Figure 1). To taxonomically characterize the predominant bacterial and archaeal community inhabiting the rhizosphere of *D. traunsteineri*, we conducted PCR amplification and sequencing of the V3-V4 variable region of the 16S rRNA gene from the soil and root wash samples in technical duplicates. Following paired-end sequencing, a total of 240,935 reads were generated, each with a read length of 300 bp (Table 1). After rigorous processing steps, including trimming, filtering of low-quality reads, merging of forward and reverse reads, and removal of chimeric sequences, 106,648 sequences were clustered into 2350 amplicon sequence variants (ASV) with taxonomic assignments spanning 31 phyla and over 84 classes (Figure 2). In total, we identified 2349 bacterial and 1 archaeal ASVs. The prevalence of the five major bacterial phyla ranged from 38.43% to 41.33% for *Proteobacteria*, 11.09% to 13.55% for *Actinobacteriota*, 9.40% to 11.90% for *Myxococcota*, 9.72% to 11.37% for *Bacteriodota*, and from 8.72% to 9.26% for *Acidobacteriota* (Table 2). Comparisons of ASV abundances across the four samples using permutational multivariate analysis of variance (PERMANOVA) with the R package vegan revealed no significant differences (similarity p-value < 0.09), indicating a consistent sampling and sequencing approach. Encouraged by the stability of the bacterial composition in the soil surrounding the roots of *D. traunsteineri* and the bacteria adhering to the roots themselves (root wash samples), we opted to analyze and process the shotgun metagenomic sequences as a unified sample.

**Figure 1.**
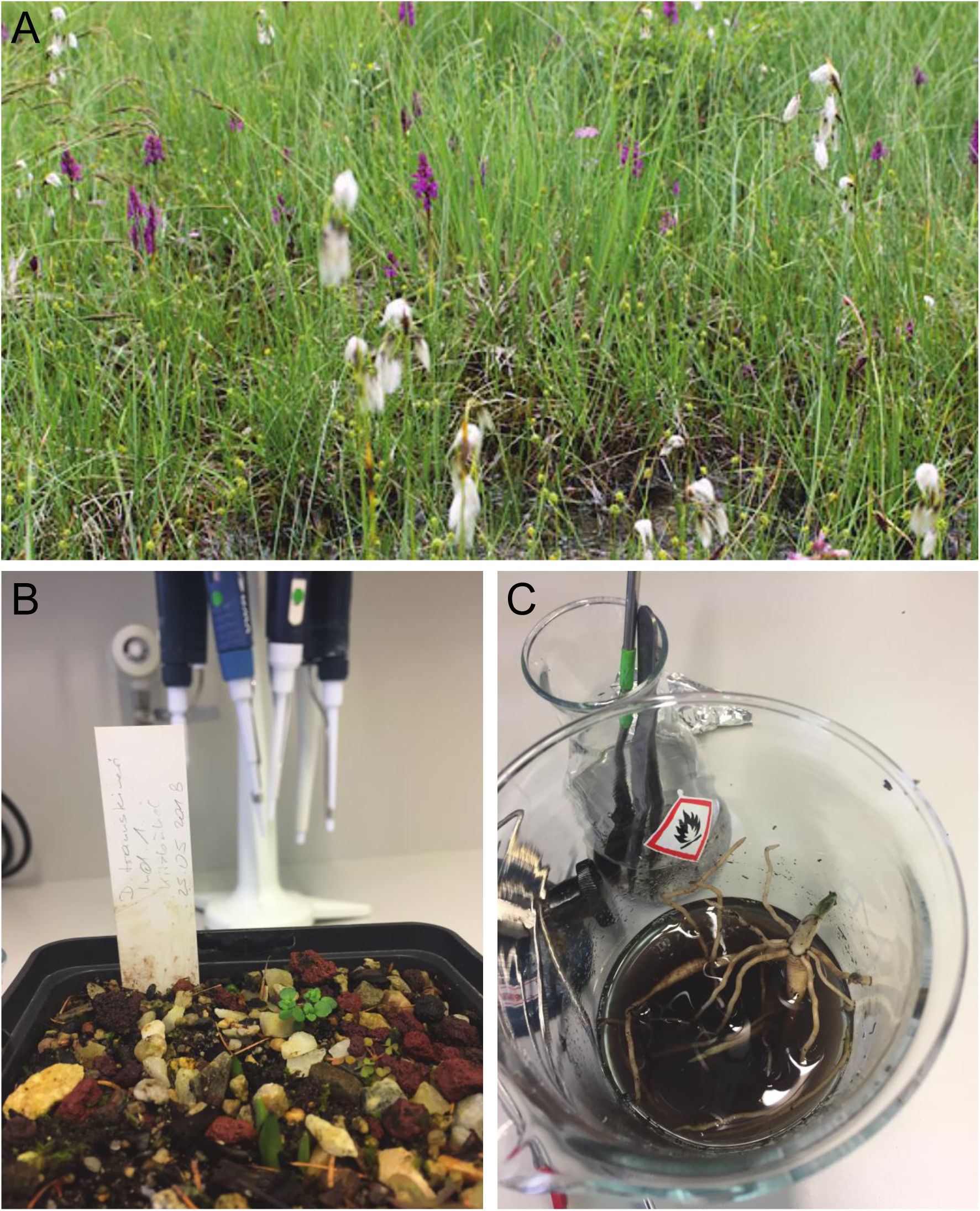
*Dactylorhiza traunsteineri* sampling. *D. traunsteineri* individuals from the Kitzbühel region (Tyrol, Austria) (**A**). *D. traunsteineri* individuum 1 from Kitzbühel in the laboratory environment (**B**). Washing of the roots with laboratory-grade water (**C**).

**Figure 2.**
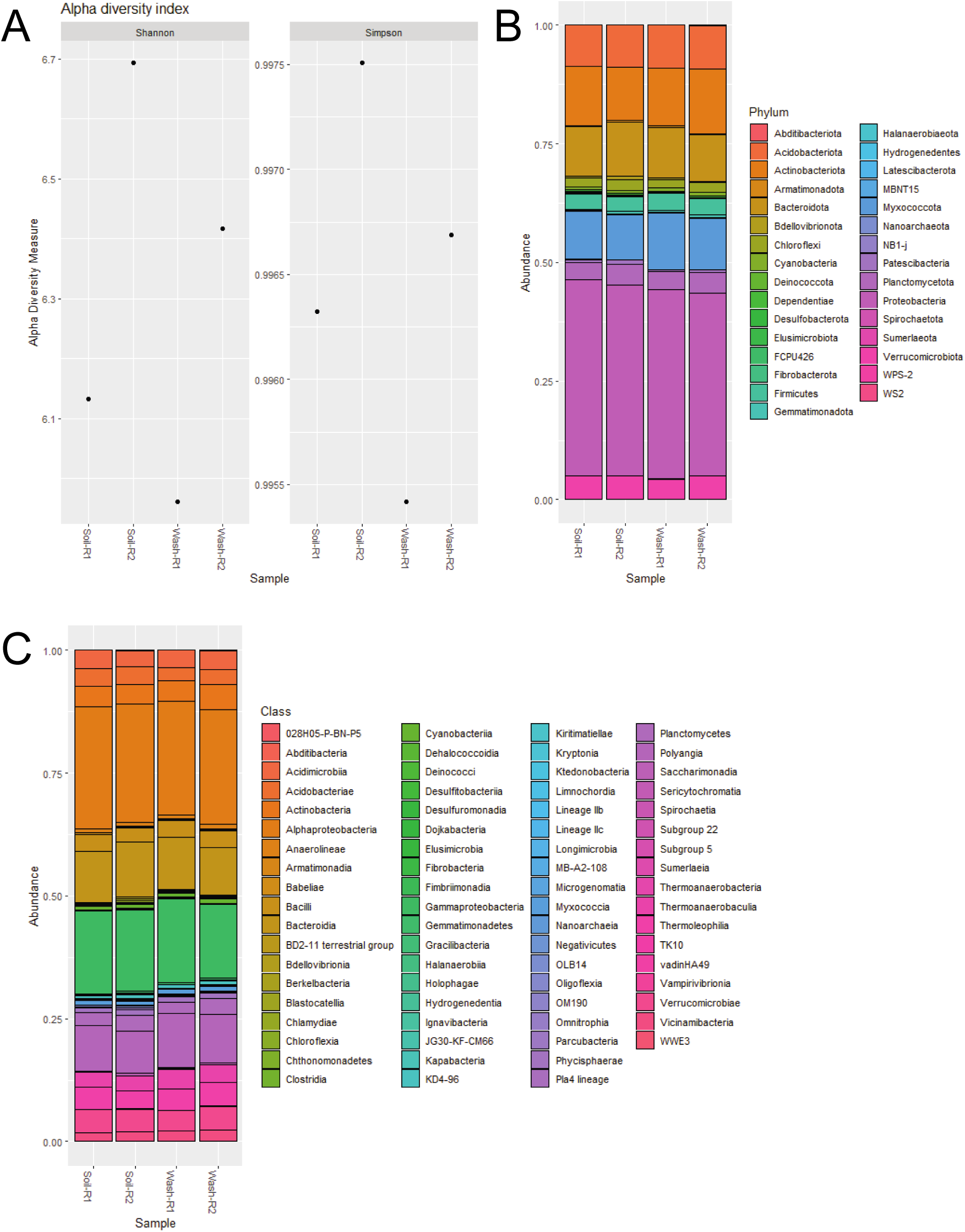
Bacterial Community Composition Analysis based on 16S rDNA sequencing. Alpha-diversity (Shannon and Simpson indexes) (**A**). Relative abundance of identified ASVs at Phylum (**B**) and Class (**C**) levels.

**Table 2.** ASVs abundance (by Phylum) (%)

|  | Soil-R1 | Soil-R2 | Wash-R1 | Wash-R2 |
| --- | --- | --- | --- | --- |
| Proteobacteria | 41.33 | 40.11 | 39.98 | 38.43 |
| Actinobacteriota | 12.41 | 11.09 | 12.20 | 13.55 |
| Myxococcota | 10.15 | 9.40 | 11.90 | 10.84 |
| Bacteroidota | 10.40 | 11.37 | 10.56 | 9.72 |
| Acidobacteriota | 8.72 | 8.85 | 9.02 | 9.26 |
| Verrucomicrobiota | 4.92 | 5.04 | 4.29 | 4.97 |
| Planctomycetota | 3.59 | 4.36 | 3.64 | 4.33 |
| Firmicutes | 3.34 | 2.88 | 3.54 | 3.40 |
| Chloroflexi | 1.82 | 2.24 | 1.87 | 2.03 |
| Cyanobacteria | 0.65 | 0.66 | 0.70 | 0.84 |
| Patescibacteria | 0.54 | 1.02 | 0.40 | 0.66 |
| Gemmatimonadota | 0.24 | 0.57 | 0.41 | 0.48 |
| Bdellovibrionota | 0.41 | 0.70 | 0.27 | 0.28 |
| Armatimonadota | 0.22 | 0.48 | 0.41 | 0.29 |
| Dependentiae | 0.35 | 0.34 | 0.17 | 0.31 |
| Desulfobacterota | 0.31 | 0.15 | 0.09 | 0.14 |
| Latescibacterota | 0.11 | 0.13 | 0.12 | 0.09 |
| FCPU4260 | 0.07 | 0.12 | 0.12 | 0.07 |
| Elusimicrobiota | 0.12 | 0.09 | 0.08 | 0.06 |
| MBNT15 | 0 | 0.10 | 0.13 | 0.07 |
| Fibrobacterota | 0.01 | 0.13 | 0.04 | 0.01 |
| Spirochaetota | 0.03 | 0.01 | 0.12 | 0.04 |
| NB1-j | 0.05 | 0.02 | 0.04 | 0.02 |
| WPS-2 | 0.09 | 0.01 | 0 | 0 |
| Sumerlaeota | 0.03 | 0.03 | 0 | 0.02 |
| Nanoarchaeota | 0.05 | 0 | 0 | 0 |
| Abditibacteriota | 0 | 0.01 | 0.02 | 0.02 |
| WS2 | 0 | 0.01 | 0 | 0.04 |
| Hydrogenedentes | 0 | 0.02 | 0 | 0.01 |
| Deinococcota | 0 | 0.01 | 0 | 0 |
| Halanaerobiaeota | 0 | 0.01 | 0 | 0 |

### Metagenome assembled genomes (MAGs) generation

To investigate the functional properties of the *D. traunsteineri* rhizobiome, we proceeded to shotgun metagenomic sequencing using short-read technology. We generated 33,043,213 paired-end metagenomic reads from the soil and root wash samples, totalling over 9 Gbp, with each sample ranging from over 1.8 to over 4.9 Gbp (Table 3). After in silico pruning of adapter sequences and low-quality reads, 31,541,795 reads were retained, representing over 95% of the initial dataset. *De novo* metagenomic assembly on the combined datasets using MEGAHIT (Li et al., 2015, Li et al., 2016) resulted in a total of 859,083 contigs, totalling over 7.5 Gbp (Table 4).

**Table 3.** Sequencing statistics (Shotgun sequencing)

| Samples | SRA accession no. | No. of raw sequence reads | No. of reads that passed quality control |
| --- | --- | --- | --- |
| Soil | SRR29434313 | 6,252,114 | 6,050,295 |
| Wash | SRR29434312 | 10,152,392 | 9,850,373 |

**Table 4.**
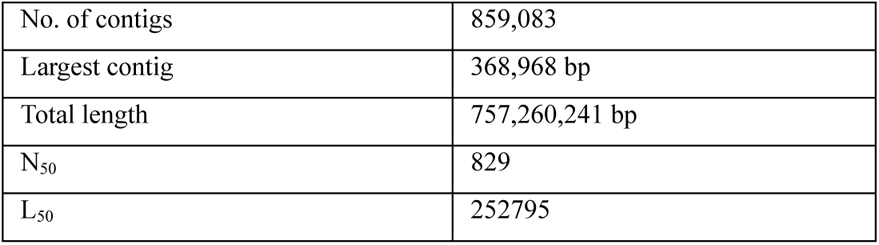
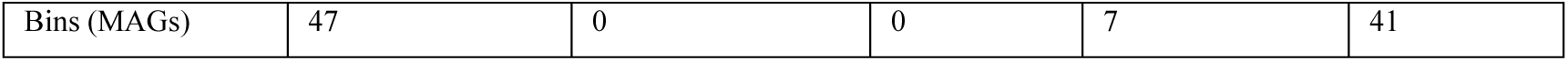
Metagenomic assembly statistics.

The obtained contigs were binned using Autometa (Miller et al., 2019), resulting in 37 predicted MAGs, as well as a group of unclustered contigs, which was further binned using MetaBAT2 (Kang et al., 2015), thus resulting in 10 additional MAGs (Figure 3). Overall, from the 248,298 contigs that were binned as Bacteria, 44,138 contigs could be grouped into 47 MAGs, from which 36 MAGs could successfully be reassembled using SPAdes (Bankevich et al., 2012, Nurk et al., 2017). These MAGs represent 359 Mbp of assembled reads, while 14.86% of all reads mapped back to the sequences contained in the combined bacterial MAGs (Table 5). MAG size ranges from 361 kbp to 45.81 Gbp, with N50 ranging from 0.9 kbp to 221 kbp (Table 6). The average predicted completeness of the MAGs stands at 33.64%, with a mean coverage spanning from 2.11 to 25.00x. Following a quality assessment, 13 initially generated MAGs were excluded due to either a contamination score exceeding 50% or completeness falling below 10%. The taxonomic distribution of the top-represented groups revealed that 28 MAGs were classified as *Proteobacteria*, eight as *Acidobacteria*, and eight as *Bacteroidetes* (Table 7). Notably, our assembly efforts yielded one high-quality draft genome (MAG_Dt_26), satisfying the stringent Minimum Information about a Metagenome-Assembled Genome (MIMAG) criteria, boasting less than 5% contamination and over 90% completeness, as well as three medium-quality draft genomes (MAG_Dt_01, MAG_Dt_03, MAG_Dt_23), with estimated completeness above 50% with less than 10% estimated contamination (Bowers et al., 2017).

**Figure 3.**
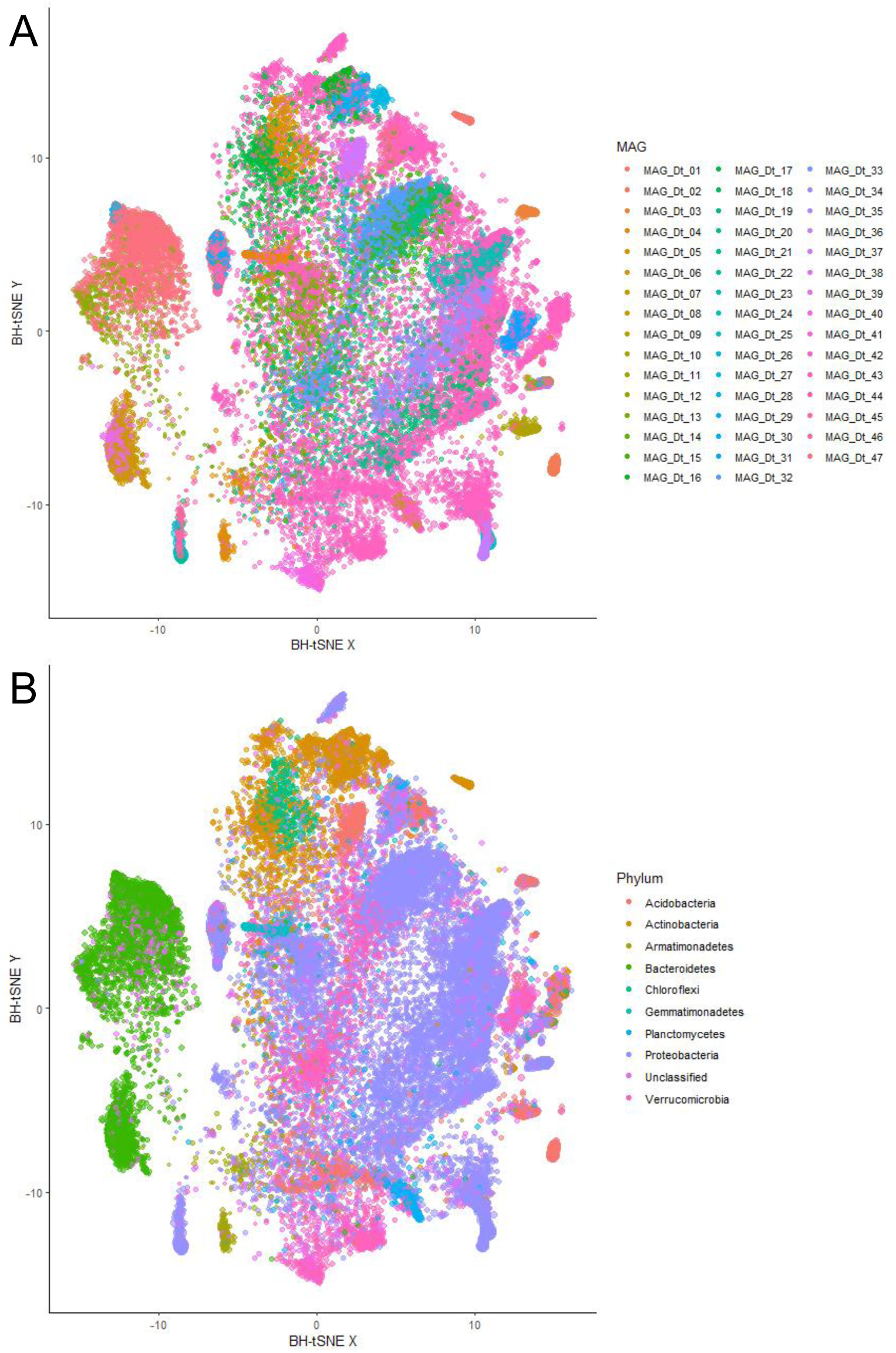
Barnes-Hut t-SNE representation of MAG contigs. BH-t-SNE representation of MAG contigs coloured by MAG (**A**) and taxonomic assignment (**B**).

**Table 5.**
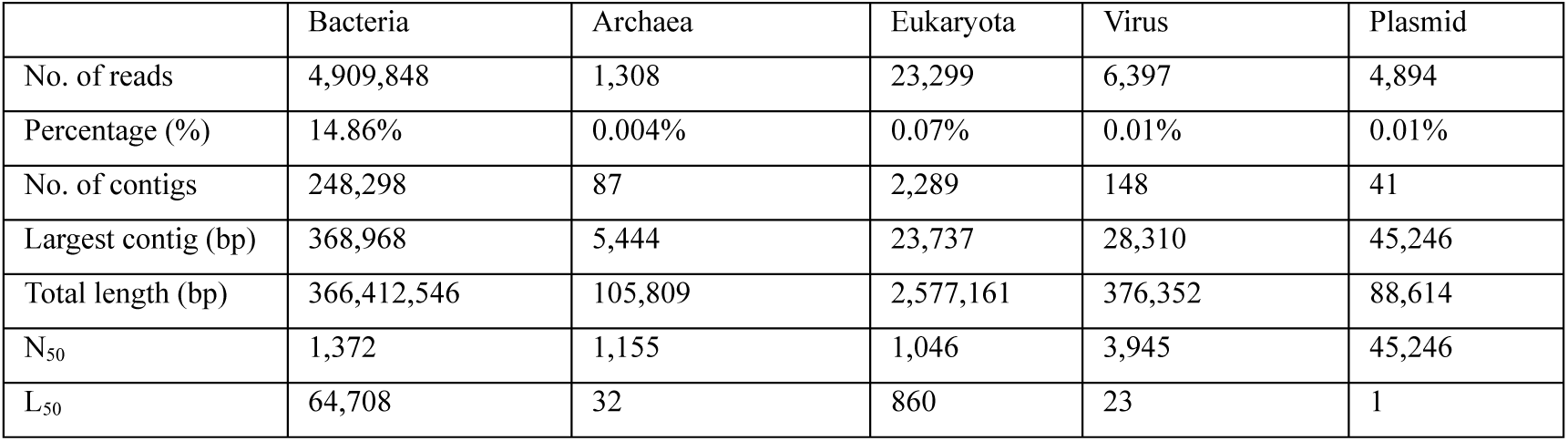
Metagenomic binning statistics.

### Gene prediction and secondary metabolite potential

Gene prediction was performed using PRODIGAL (Hyatt et al., 2010) for each MAG, and the resulting predicted proteins were annotated using the KEGG (Kanehisa and Goto, 2000, Kanehisa et al., 2023) and PANNZER2 (Toronen et al., 2018) databases for metabolic pathway enrichment (Table 8). Furthermore, each MAG was mined for SM BSGs using the nf-core funcscan pipeline (Ewels et al., 2020). A total of 1,741 BGCs were predicted from the *D. traunsteineri* rhizosphere bacterial MAGs (Figure 4; Table 9).

**Figure 4.**
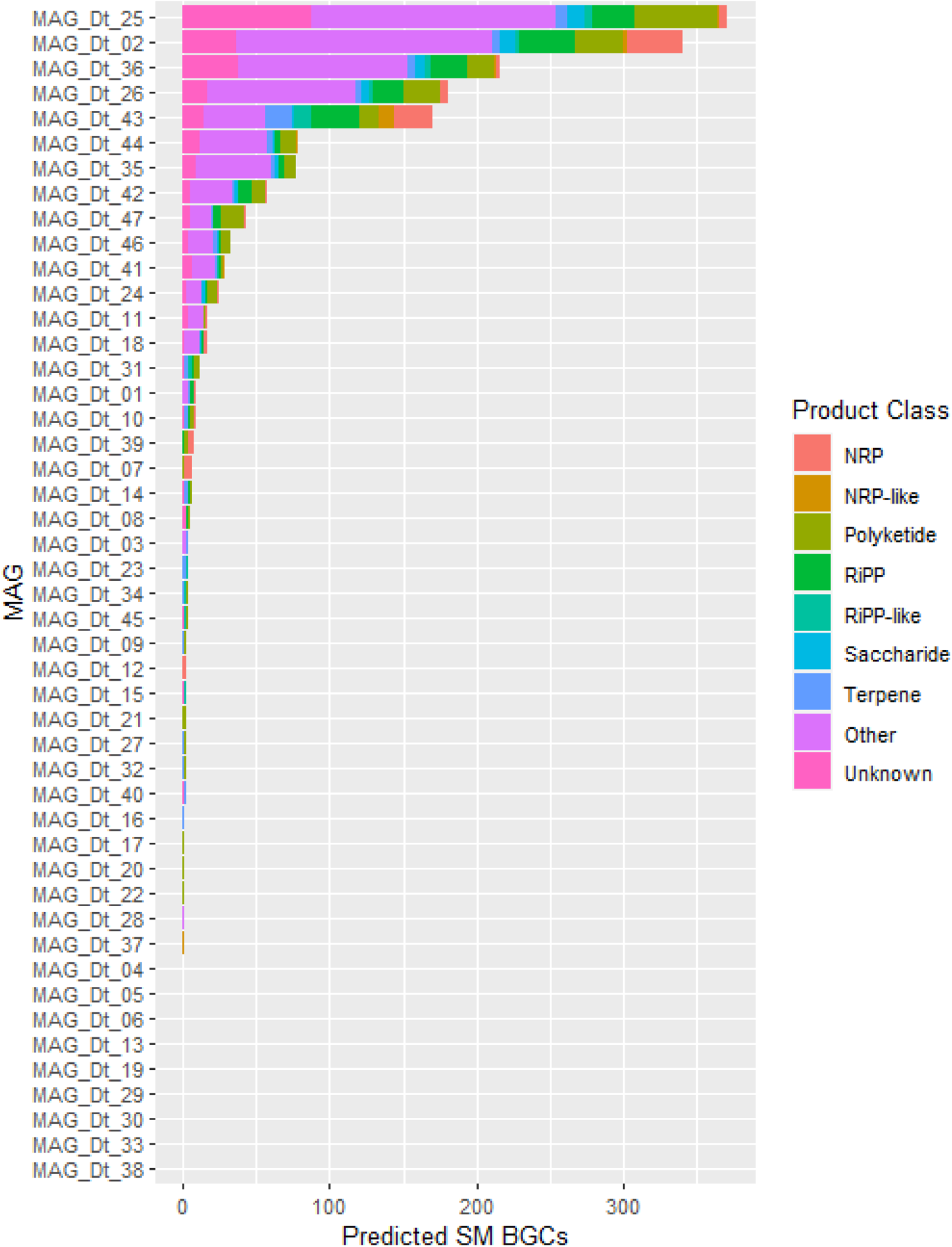
Overview of number of predicted SM BGCs per MAG and their product classes.

In MAG_Dt_26 (*Proteobacteria*; completeness of 92.03%; contamination of 2.53%), a total of 181 SM BGCs were predicted. These included six nonribosomal peptide (NRP) clusters, 24 polyketide clusters, 22 ribosomally synthesized and post-translationally modified peptide (RiPP) clusters, two RiPP-like clusters, five saccharide clusters, as well as four terpene clusters (Supp. Table 2). Additionally, 101 gene clusters were categorized as “Other”, potentially representing multiple product types and rarer SM classes, while 17 clusters were classified as “Unknown” (Supp. Table 2). From MAG_Dt_01 (*Actinobacteria*; completeness of 80.24%; contamination of 4.58%), a total of nine SM BGCs were predicted, representing one NRP cluster, one polyketide cluster, two RiPP clusters, two terpene clusters, as well as three “Other” product class clusters (Supp. Table 2). From MAG_Dt_03 (*Acidobacteria*; completeness of 65.94%; contamination of 3.99%), four SM BGCs were predicted, representing one NRP cluster, one terpene cluster, and two “Other” product class clusters (Supp. Table 2). From MAG_Dt_23 (*Proteobacteria*; completeness of 61.01%; contamination of 2.68%), four SM BGCs were predicted, representing two RiPP-like and two terpene clusters (Supp. Table 2).

## DISCUSSION

Through comprehensive 16S rDNA profiling, a total of 2,349 bacterial and one archaeal ASVs were identified within the rhizosphere of *D. traunsteineri*. The predominant bacterial phyla detected included *Proteobacteria*, *Actinobacteria*, *Myxococcota*, *Bacteroidota*, and *Acidobacteria*, which collectively accounted for 95% of the sequences identified. Importantly, there were no significant differences in the bacterial community composition between the soil and the root-adhering bacteria associated with *D. traunsteineri*. This consistency underscores the robustness and reproducibility of our sample preparation and sequencing methodologies. Moreover, in our review of existing literature on orchid RAB ecology, we validated that our approach successfully recovered representatives from all phyla previously reported in orchid rhizospheres (Kaur and Sharma, 2021). Notably, the scientific documentation of orchid RAB through culture-independent methods is relatively scarce. Previous studies have predominantly utilized conventional Sanger sequencing methods, while others employing high-throughput sequencing technologies have primarily focused on 16S rDNA profiling of bacterial communities directly extracted from root samples. Nevertheless, the field of orchid RAB research is undergoing a transformative shift with the advent of innovative methodologies. In this context, we introduce a pioneering approach that leverages shotgun metagenomics, followed by the reconstruction of MAGs. This cutting-edge technique promises to offer deeper insights into the complex ecosystem of the orchid rhizobiome, providing unparalleled understanding of its microbial diversity and functional potential. Utilizing shotgun sequencing and *de novo* metagenomic assembly, we successfully obtained 47 bacterial MAGs from the rhizobiome of *D. traunsteineri.* Our detailed analysis focused on four high-quality MAGs: MAG_Dt_01, MAG_Dt_03, MAG_Dt_23, and MAG_Dt_26. Specifically, MAG_Dt_01 was taxonomically assigned to *Actinobacteria bacterium* (taxid 1883427), MAG_Dt_03 to *Acidobacteria bacterium* (taxid 1978231), and MAG_Dt_23 and MAG_Dt_26 to *Dongia* sp. URHE0060 (taxid 1380391) and *Steroidobacter* (taxid 469322), respectively, both belonging to the phylum *Proteobacteria*.

The phyla *Proteobacteria* and *Actinobacteria*, generally enriched in the rhizosphere, alongside *Acidobacteria*, typically enriched in bulk soil, play pivotal roles in soil carbon and nitrogen cycles as well as the decomposition of organic matter (Eichorst et al., 2018, Song et al., 2021, Ling et al., 2022). The *Actinobacteria* phylum represents one of the largest and most diverse bacterial group, inhabiting soil, water and plant tissue (Diab et al., 2024, Boubekri et al., 2022), and are prolific producers of SMs including well-known antibiotics such as streptomycin, tetracycline, erythromycin, and vancomycin (Berdy, 2005). Beyond antibiotics, they produce a wide array of SMs with potential applications as biopesticides (Barka et al., 2016). For example, polyketides, such as macrolides, exhibit antifungal activity, while non-ribosomal peptides (NRPs), like cyclopeptides, demonstrate insecticidal properties (Gaynor and Mankin, 2003, Quiroz-Carreno et al., 2022). The *Acidobacteria* phylum, although highly prevalent within soil ecosystems, is characterized by its challenging cultivation (Kalam et al., 2020). This phylum fulfills crucial ecological functions through its significant involvement in carbon, nitrogen, and sulphur biogeochemical processes (Ward et al., 2009, Eichorst et al., 2018, Hausmann et al., 2018). Furthermore, *Acidobacteria* possess genes conducive to survival and competitive colonization within the rhizosphere, thereby facilitating the formation of symbiotic relationships with plants (Kalam et al., 2020, Liu et al., 2024). Members of the *Proteobacteria* phylum, such as the *Dongia* and *Steroidobacter* genera, have been frequently isolated from rhizosphere soil and display the potential to degrade complex organic compounds, including various polysaccharides (Ikenaga et al., 2021, Huang et al., 2019, Liu et al., 2024, Baik et al., 2013, Kim et al., 2016, Liu et al., 2010, Palla et al., 2022). The high-quality MAGs obtained in this study present a promising opportunity to elucidate in greater detail the roles of these phyla in plant microbiome interactions, particularly regarding their secondary metabolite potential and their impact on plant health and development.

Given that our sampling and sequencing efforts were confined to the rhizobiome of a solitary specimen of *D. traunsteineri*, caution must be exercised in generalizing the presence and prevalence of the identified MAGs across all populations of *D. traunsteineri*. However, it is important to note that the sampled individual originated from an extant population and has been maintained under conditions that closely simulate its natural habitat at the Botanical Garden of the University of Vienna. These conditions include soil composition, humidity, and temperature, which are critical for mimicking the plant’s natural environment. The controlled yet naturalistic setting in which the specimen was studied provides a valuable context for our findings. Despite the inherent limitations of a single-sample study, our research offers a foundational framework for future interaction studies between RABs and *D. traunsteineri*. This study lays the groundwork for more extensive investigations into the microbiome of this orchid species across different populations and environmental conditions. Furthermore, our findings open new avenues for exploring the complex processes involved in the adaptation, ecology, and evolution of this allopolyploid orchid. Understanding the interactions between *D. traunsteineri* and its associated microbiome can shed light on the mutualistic relationships that facilitate plant health, nutrient acquisition, and stress resilience. Such insights are crucial for conservation strategies, especially for orchids that are often sensitive to environmental changes and habitat disturbances. Additionally, the use of advanced metagenomic techniques in our study underscores the potential for discovering novel microbial taxa and functional genes that contribute to the unique ecology of orchid rhizospheres. This approach not only enhances our understanding of plant-microbe interactions but also provides a deeper comprehension of the evolutionary dynamics within the orchid family. Our study, therefore, serves as a stepping stone for future research aimed at unraveling the intricate web of interactions that sustain and influence the biodiversity and ecological success of *D. traunsteineri*.

## CONCLUSION

This study provided a comprehensive overview of the microbiome inhabiting the rhizosphere of the endangered orchid *Dactylorhiza traunsteineri*. By conducting an in-depth bacterial community composition analysis through 16S rDNA-targeted sequencing, we identified *Proteobacteria*, *Actinobacteria*, *Myxococcota*, *Bacteroidota*, and *Acidobacteria* as the most abundant phyla within the *D. traunsteineri* rhizosphere. This foundational data highlights the complexity and diversity of the microbial communities associated with this orchid species. Our investigation extended beyond community profiling to include deep shotgun metagenomics sequencing and subsequent *de novo* metagenome assembly. This approach enabled us to extract high-quality metagenome-assembled genomes (MAGs), providing a rich resource for further functional analysis. Each MAG was meticulously analyzed for metabolic pathway enrichment and secondary metabolite (SM) biosynthetic gene clusters (BGCs), revealing a previously unexplored niche of the rhizobiome. These analyses uncovered potential metabolic capabilities and biosynthetic potentials that are integral to the rhizosphere’s ecological dynamics.

The insights gained from this study not only enhance our understanding of the microbial diversity within the *D. traunsteineri* rhizosphere but also lay the groundwork for future research aimed at unravelling the intricate interconnections between the rhizobiome and its host. The high-quality MAGs we have made available serve as a valuable resource for further exploration of microbial functions and their contributions to the health, adaptation, and survival of *D. traunsteineri*. Moreover, our findings underscore the importance of employing advanced metagenomic techniques to study plant-associated microbiomes, particularly for endangered species. By elucidating the functional roles of key microbial players, we can develop more effective conservation strategies and foster a deeper understanding of the symbiotic relationships that underpin plant resilience and ecological success. In conclusion, our study represents a significant step forward in orchid microbiome research, providing novel insights and a robust platform for future investigations into the dynamic interactions within the rhizosphere of *D. traunsteineri*. These efforts are crucial for advancing our knowledge of plant-microbe interactions and supporting the conservation of this and other endangered orchid species.

## Software

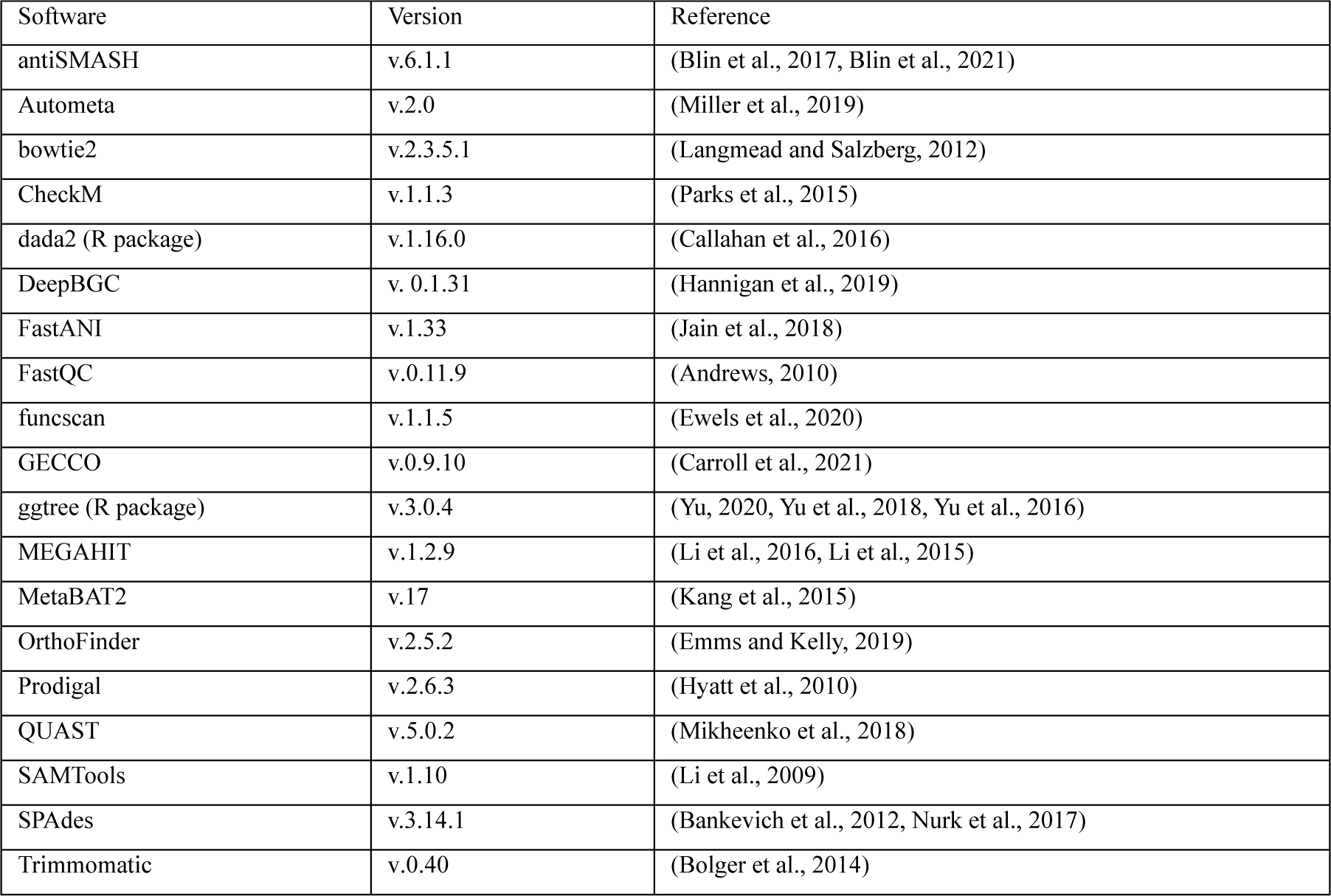

## Abbreviations

ANI: average nucleotide identity
ASV: amplicon sequence variant
BGC: biosynthetic gene cluster
MAG: metagenome assembled genome
MIMAG: Minimum Information about a Metagenome-Assembled Genome
OTU: operational taxonomic unit
PERMANOVA: permutational multivariate analysis of variance
PGPR: plant growth promoting Rhizobacteria
RAB: root associated bacteria
SM: secondary metabolite

## Authors’ contribution

GAV, CZ, DFS and OV conceptualized and designed the study. GAV and LZ generated the datasets. GAV and JC performed the data analysis. GAV and JC generated the figures and drafted the manuscript. ARMA and RLM revised and contributed to the manuscript. All authors read and approved the final manuscript.

## Acknowledgements

We would like to thank David Pressler and the Botanical Garden of the University of Vienna for maintaining the plant in cultivation, and the Tyrol County administration for issuing necessary permits.

## Availability of data and materials

The datasets supporting the conclusions of this article are publicly available in the National Center for Biotechnology Information (NCBI) repository under BioProject PRJNA1124010.

## Ethics approval and consent to participate

Not applicable.

## Consent for publication

Not applicable.

## Corresponding author

Correspondence to Julien Charest

## Supplementary Information

**Supplementary Figure 1.**
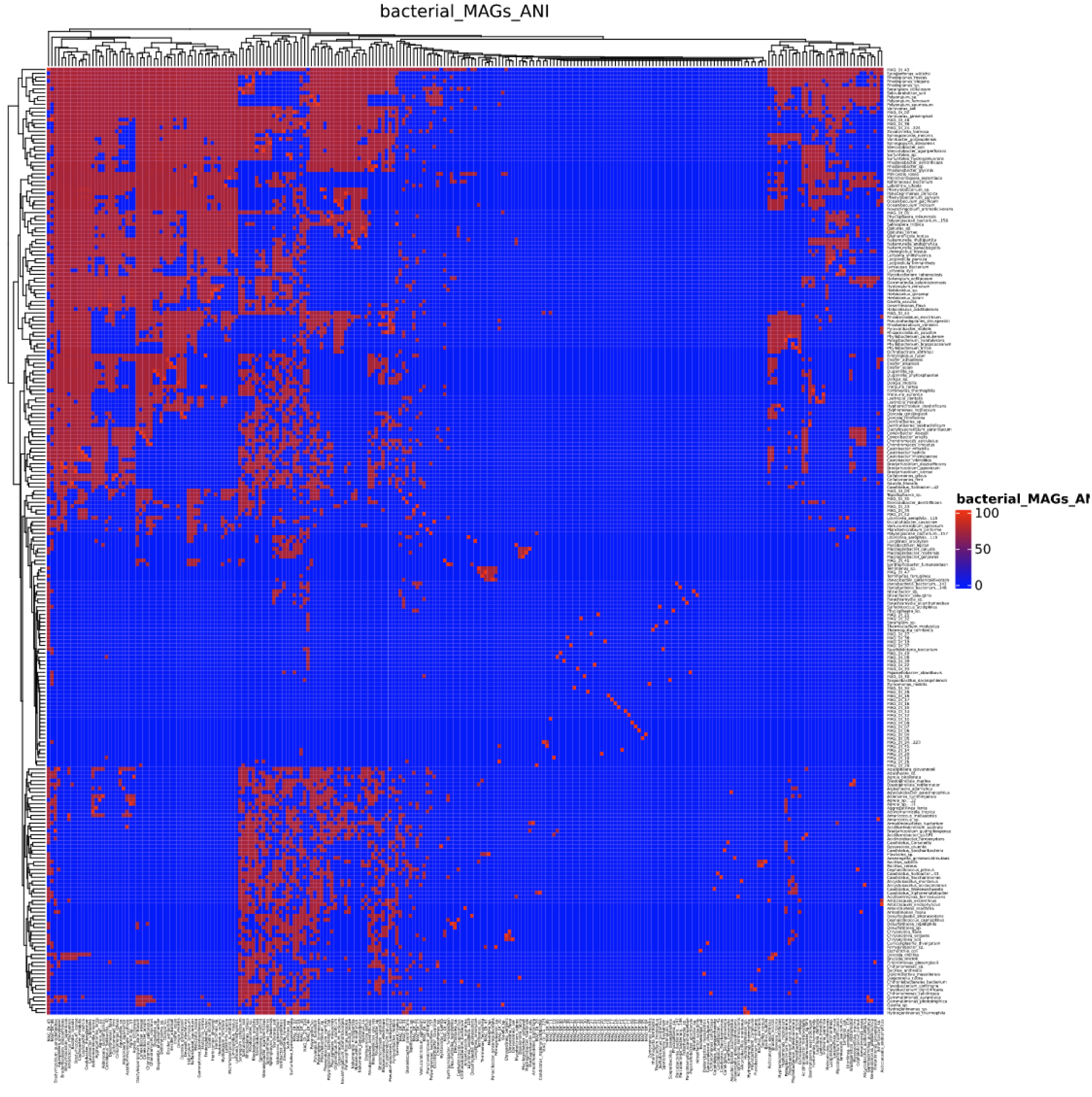
ANI of MAGs and reference organisms.

**Supplementary Table 1.**
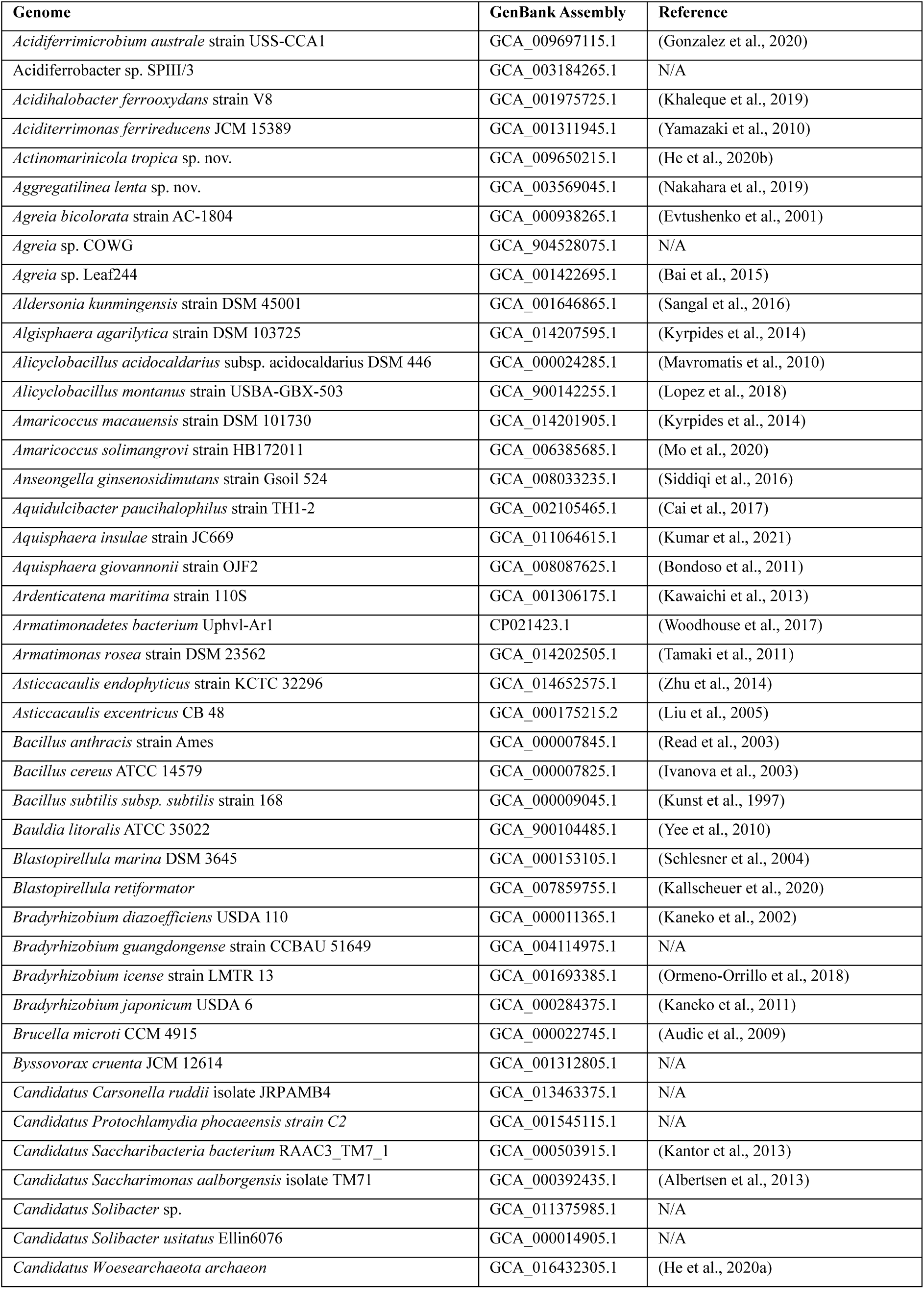
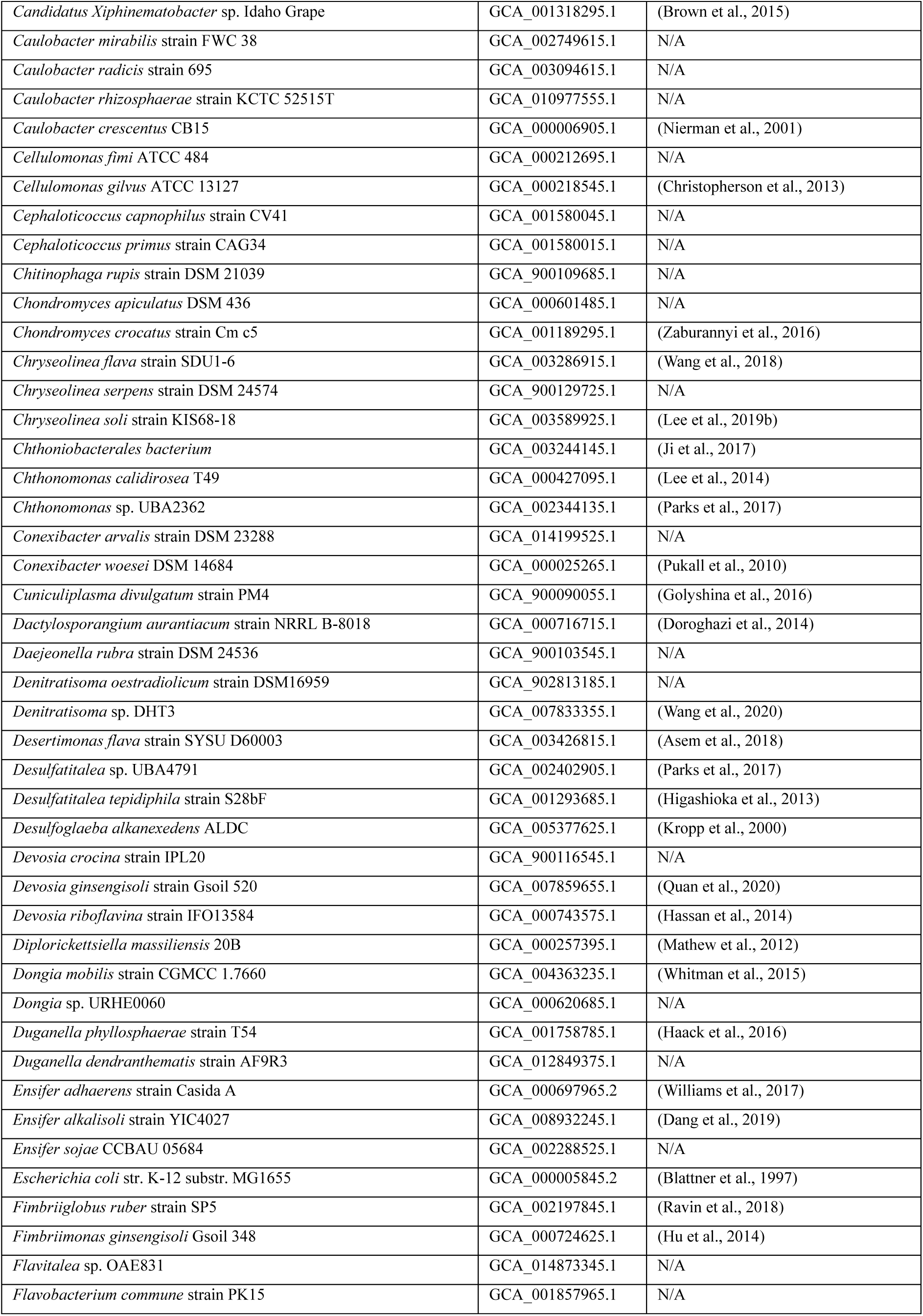
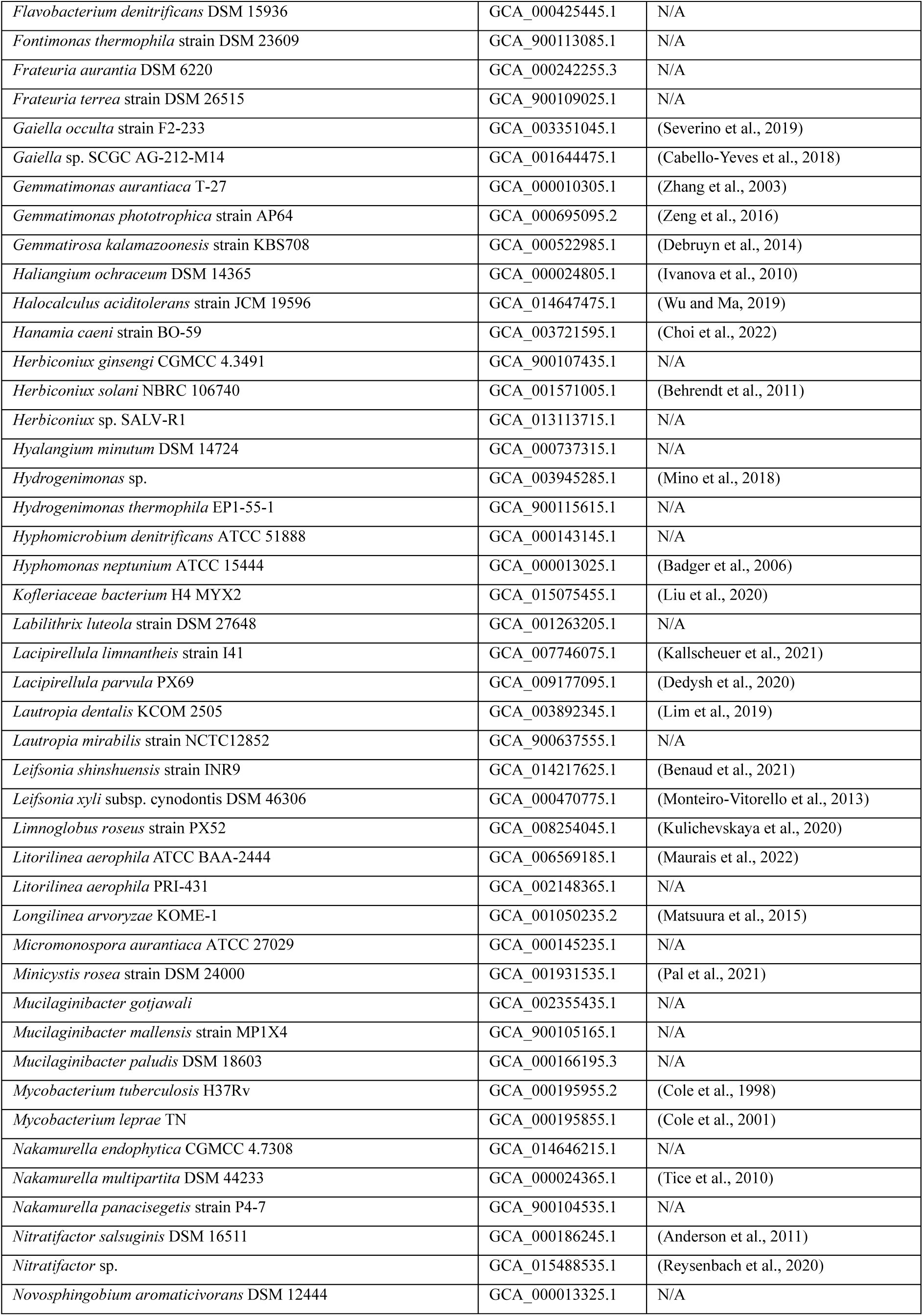
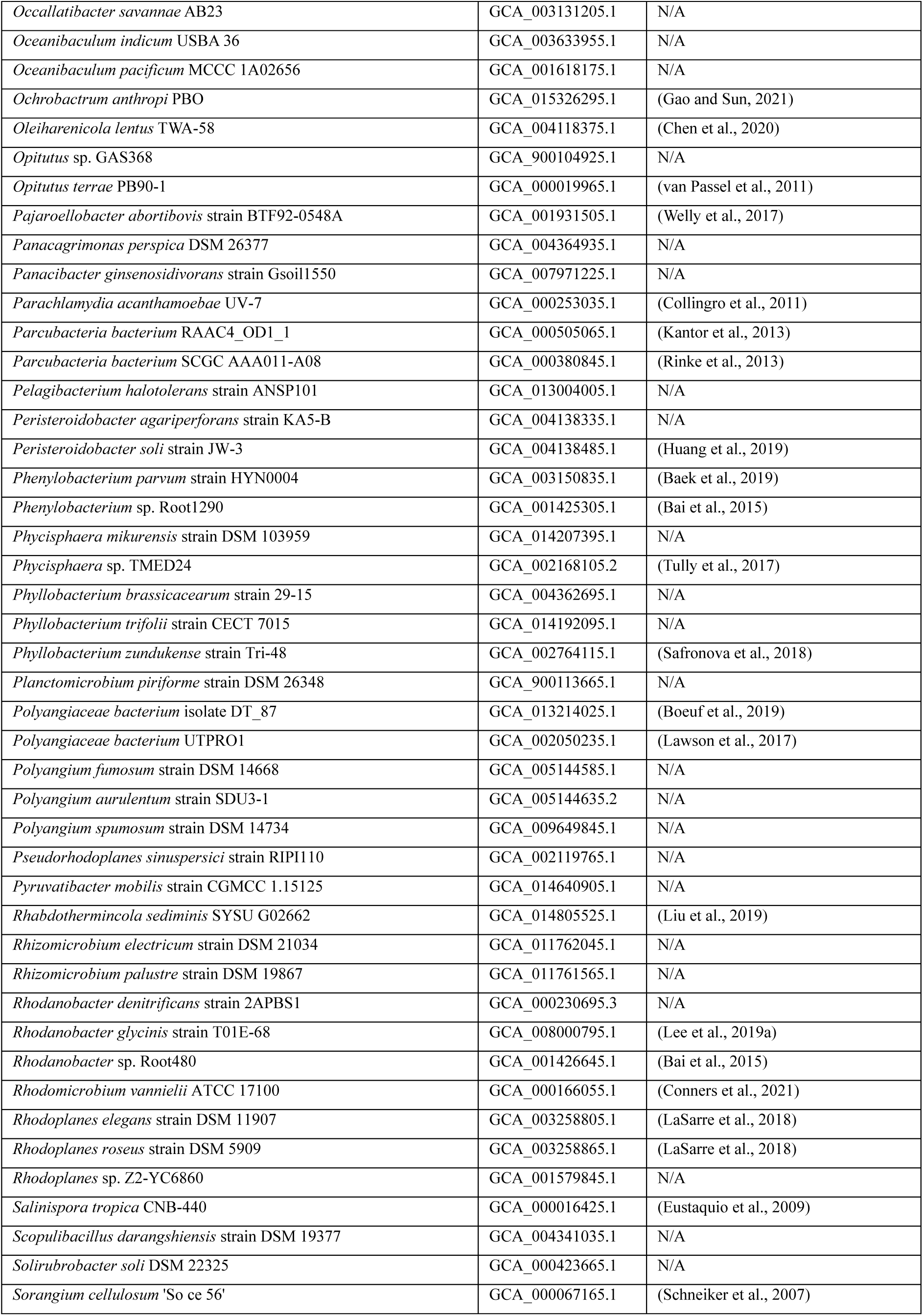
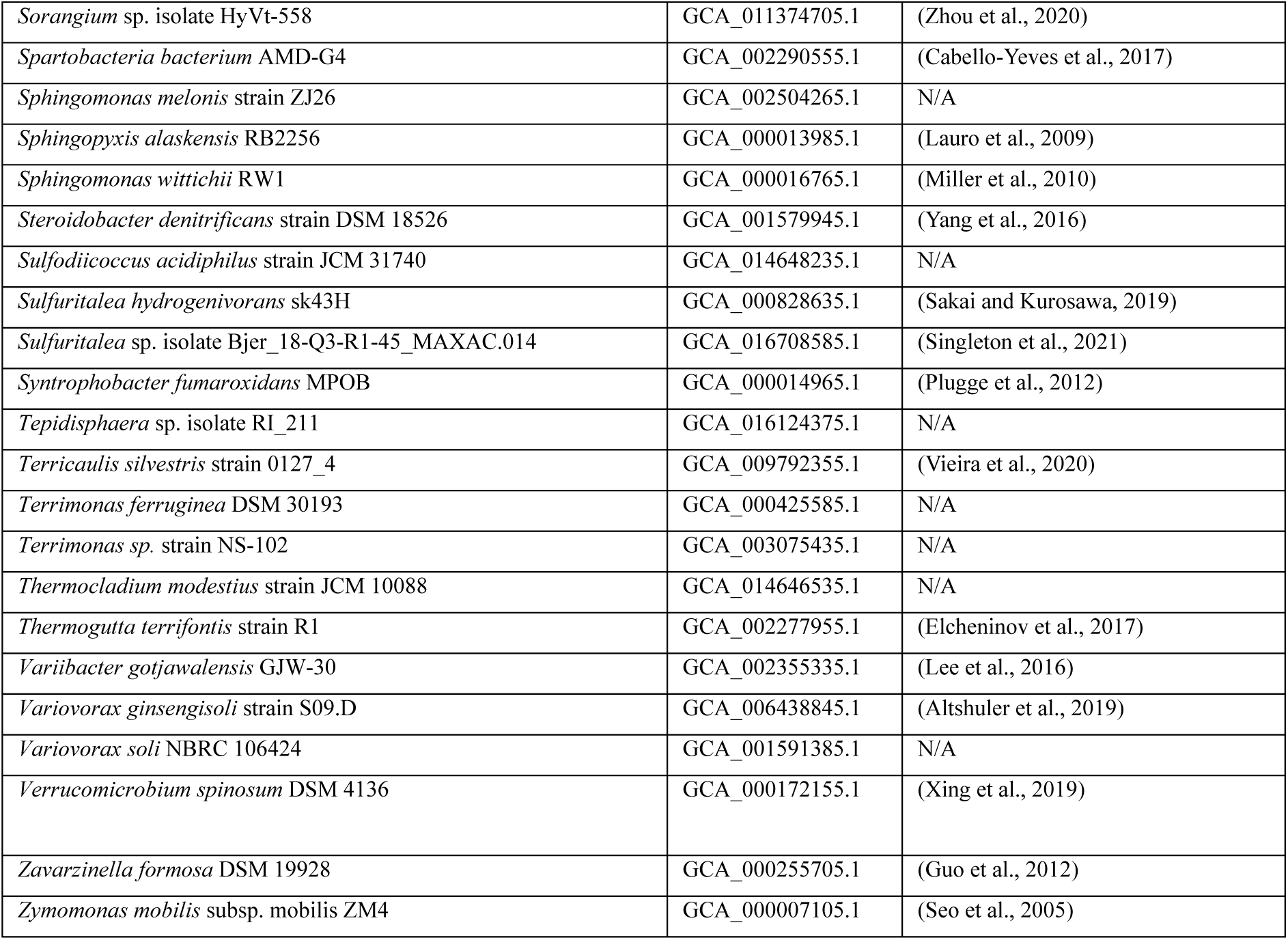
Genomes used for comparative analysis.

## REFERENCES

Andrews, S. 2010. FastQC: A Quality Control Tool for High Throughput Sequence Data [Online]. Available: http://www.bioinformatics.babraham.ac.uk/projects/fastqc/ [Accessed].

Backer, R., Rokem, J. S., Ilangumaran, G., Lamont, J., Praslickova, D., Ricci, E., Subramanian, S. & Smith, D. L. 2018. Plant Growth-Promoting Rhizobacteria: Context, Mechanisms of Action, and Roadmap to Commercialization of Biostimulants for Sustainable Agriculture. Front Plant Sci, 9, 1473.

Baik, K. S., Hwang, Y. M., Choi, J. S., Kwon, J. & Seong, C. N. 2013. Dongia rigui sp. nov., isolated from freshwater of a large wetland in Korea. Antonie Van Leeuwenhoek, 104, 1143–50.

Bankevich, A., Nurk, S., Antipov, D., Gurevich, A. A., Dvorkin, M., Kulikov, A. S., Lesin, V. M., Nikolenko, S. I., Pham, S., Prjibelski, A. D., Pyshkin, A. V., Sirotkin, A. V., Vyahhi, N., Tesler, G., Alekseyev, M. A. & Pevzner, P. A. 2012. SPAdes: a new genome assembly algorithm and its applications to single-cell sequencing. J Comput Biol, 19, 455–77.

Barka, E. A., Vatsa, P., Sanchez, L., Gaveau-Vaillant, N., Jacquard, C., Meier-Kolthoff, J. P., Klenk, H. P., Clement, C., Ouhdouch, Y. & Van Wezel, G. P. 2016. Taxonomy, Physiology, and Natural Products of Actinobacteria. Microbiol Mol Biol Rev, 80, 1–43.

Bartók, A., Brădeanu, A., Dulugeac, R., Bobocea, M.-M. & Şesan, T. E. 2018. Dactylorhiza traunsteineri (Saut. ex Reichenb.) Soó Románia flórájában. Kitaibelia, 23, 15–24.

Baumann, H. & Künkele, S. 1988. *Die Orchideen Europas*, Stuttgart, Kosmos Naturführe.

Berdy, J. 2005. Bioactive microbial metabolites. J Antibiot (Tokyo), 58, 1–26.

Blin, K., Shaw, S., Kloosterman, A. M., Charlop-Powers, Z., Van Wezel, G. P., Medema, M. H. & Weber, T. 2021. antiSMASH 6.0: improving cluster detection and comparison capabilities. Nucleic Acids Res, 49, W29–W35.

Blin, K., Wolf, T., Chevrette, M. G., Lu, X., Schwalen, C. J., Kautsar, S. A., Suarez Duran, H. G., De Los Santos, E. L. C., Kim, H. U., Nave, M., Dickschat, J. S., Mitchell, D. A., Shelest, E., Breitling, R., Takano, E., Lee, S. Y., Weber, T. & Medema, M. H. 2017. antiSMASH 4.0-improvements in chemistry prediction and gene cluster boundary identification. Nucleic Acids Res, 45, W36–W41.

Blinova, I. V. & Uotila, P. 2012. Dactylorhiza traunsteineri (Orchidaceae) in Murmansk Region (Russia). Memoranda Societatis pro Fauna et Flora Fennica, 88, 67–79.

Bloemberg, G. V. & Lugtenberg, B. J. 2001. Molecular basis of plant growth promotion and biocontrol by rhizobacteria. Curr Opin Plant Biol, 4, 343–50.

Bolger, A. M., Lohse, M. & Usadel, B. 2014. Trimmomatic: a flexible trimmer for Illumina sequence data. Bioinformatics, 30, 2114–20.

Boubekri, K., Soumare, A., Mardad, I., Lyamlouli, K., Ouhdouch, Y., Hafidi, M. & Kouisni, L. 2022. Multifunctional role of Actinobacteria in agricultural production sustainability: A review. Microbiol Res, 261, 127059.

Bowers, R. M., Kyrpides, N. C., Stepanauskas, R., Harmon-Smith, M., Doud, D., Reddy, T. B. K., Schulz, F., Jarett, J., Rivers, A. R., Eloe-Fadrosh, E. A., Tringe, S. G., Ivanova, N. N., Copeland, A., Clum, A., Becraft, E. D., Malmstrom, R. R., Birren, B., Podar, M., Bork, P., Weinstock, G. M., Garrity, G. M., Dodsworth, J. A., Yooseph, S., Sutton, G., Glockner, F. O., Gilbert, J. A., Nelson, W. C., Hallam, S. J., Jungbluth, S. P., Ettema, T. J. G., Tighe, S., Konstantinidis, K. T., Liu, W. T., Baker, B. J., Rattei, T., Eisen, J. A., Hedlund, B., Mcmahon, K. D., Fierer, N., Knight, R., Finn, R., Cochrane, G., Karsch- Mizrachi, I., Tyson, G. W., Rinke, C., GENOME Standards, C., Lapidus, A., Meyer, F., Yilmaz, P., Parks, D. H., Eren, A. M., Schriml, L., Banfield, J. F., Hugenholtz, P. & Woyke, T. 2017. Minimum information about a single amplified genome (MISAG) and a metagenome-assembled genome (MIMAG) of bacteria and archaea. Nat Biotechnol, 35, 725–731.

Brandrud, M. K., Baar, J., Lorenzo, M. T., Athanasiadis, A., Bateman, R. M., Chase, M. W., Hedren, M. & Paun, O. 2020. Phylogenomic Relationships of Diploids and the Origins of Allotetraploids in Dactylorhiza (Orchidaceae). Syst Biol, 69, 91–109.

Bulgarelli, D., Schlaeppi, K., Spaepen, S., Ver Loren Van Themaat, E. & Schulze-Lefert, P. 2013. Structure and functions of the bacterial microbiota of plants. Annu Rev Plant Biol, 64, 807–38.

Callahan, B. J., Mcmurdie, P. J., Rosen, M. J., Han, A. W., Johnson, A. J. & Holmes, S. P. 2016. DADA2: High-resolution sample inference from Illumina amplicon data. Nat Methods, 13, 581–3.

Carroll, L. M., Larralde, M., Fleck, J. S., Ponnudurai, R., Milanese, A., Cappio, E. & Zeller, G. 2021. Accurate de novo identification of biosynthetic gene clusters with GECCO. bioRxiv, 2021.05.03.442509.

Delforge, P. 2001. Guide des Orchidées d Éurope, d Áfrique du Nord et du Proche-Orient, Lausanne & Paris, Delachaux et Niestlé. Diab, M. K., Mead, H. M., Ahmed Khedr, M. M., Abu-Elsaoud, A. M. & El-Shatoury, S. A. 2024. Actinomycetes are a natural resource for sustainable pest control and safeguarding agriculture. Arch Microbiol, 206, 268.

Dobbelaere, S., Vanderleyden, J. & Okon, Y. 2003. Plant Growth-Promoting Effects of Diazotrophs in the Rhizosphere. Critical Reviews in Plant Sciences, 22, 107–149.

Eichorst, S. A., Trojan, D., Roux, S., Herbold, C., Rattei, T. & Woebken, D. 2018. Genomic insights into the Acidobacteria reveal strategies for their success in terrestrial environments. Environ Microbiol, 20, 1041–1063.

Emelianova, K., Hawranek, A.-S., Eriksson, M. C., Wolfe, T. M. & Paun, O. 2024. Ecological divergence is marked by species specific plastic gene expression and distinct rhizosphere among sibling allopolyploid marsh orchids (Dactylorhiza). *bioRxiv*, 2024.04.24.590886.

Emms, D. M. & Kelly, S. 2019. OrthoFinder: phylogenetic orthology inference for comparative genomics. Genome Biol, 20, 238.

Ewels, P. A., Peltzer, A., Fillinger, S., Patel, H., Alneberg, J., Wilm, A., Garcia, M. U., Di Tommaso, P. & Nahnsen, S. 2020. The nf-core framework for community-curated bioinformatics pipelines. Nat Biotechnol, 38, 276–278.

Fontaine, L., Pin, L., Savio, D., Friberg, N., Kirschner, A. K. T., Farnleitner, A. H. & Eiler, A. 2023. Bacterial bioindicators enable biological status classification along the continental Danube river. Commun Biol, 6, 862.

Fouillaud, M. & Dufosse, L. 2022. Microbial Secondary Metabolism and Biotechnology. Microorganisms, 10.

Gaynor, M. & Mankin, A. S. 2003. Macrolide antibiotics: binding site, mechanism of action, resistance. Curr Top Med Chem, 3, 949–61.

Givnish, T. J., Spalink, D., Ames, M., Lyon, S. P., Hunter, S. J., Zuluaga, A., Doucette, A., Caro, G. G., Mcdaniel, J., Clements, M. A., Arroyo, M. T. K., Endara, L., Kriebel, R., Williams, N. H. & Cameron, K. M. 2016. Orchid historical biogeography, diversification, Antarctica and the paradox of orchid dispersal. Journal of Biogeography, 43, 1905–1916.

Givnish, T. J., Spalink, D., Ames, M., Lyon, S. P., Hunter, S. J., Zuluaga, A., Iles, W. J., Clements, M. A., Arroyo, M. T., Leebens-Mack, J., Endara, L., Kriebel, R., Neubig, K. M., Whitten, W. M., Williams, N. H. & Cameron, K. M. 2015. Orchid phylogenomics and multiple drivers of their extraordinary diversification. Proc Biol Sci, 282.

Glick, B. R. 1995. The Enhancement of Plant-Growth by Free-Living Bacteria. Canadian Journal of Microbiology, 41, 109–117.

Glick, B. R. 2012. Plant growth-promoting bacteria: mechanisms and applications. Scientifica (Cairo), 2012, 963401.

Gouda, S., Kerry, R. G., Das, G., Paramithiotis, S., Shin, H. S. & Patra, J. K. 2018. Revitalization of plant growth promoting rhizobacteria for sustainable development in agriculture. Microbiol Res, 206, 131–140.

Hannigan, G. D., Prihoda, D., Palicka, A., Soukup, J., Klempir, O., Rampula, L., Durcak, J., Wurst, M., Kotowski, J., Chang, D., Wang, R., Piizzi, G., Temesi, G., Hazuda, D. J., Woelk, C. H. & Bitton, D. A. 2019. A deep learning genome-mining strategy for biosynthetic gene cluster prediction. Nucleic Acids Res, 47, e110.

Hassani, M. A., Duran, P. & Hacquard, S. 2018. Microbial interactions within the plant holobiont. Microbiome, 6, 58.

Hausmann, B., Pelikan, C., Herbold, C. W., Kostlbacher, S., Albertsen, M., Eichorst, S. A., GLAVINA DEL Rio, T., Huemer, M., Nielsen, P. H., Rattei, T., Stingl, U., Tringe, S. G., Trojan, D., Wentrup, C., Woebken, D., Pester, M. & Loy, A. 2018. Peatland Acidobacteria with a dissimilatory sulfur metabolism. ISME J, 12, 1729–1742.

Huang, J. W., Hu, S. L., Cheng, X. K., Chen, D., Kong, X. K. & Jiang, J. D. 2019. Steroidobacter soli sp. nov., isolated from farmland soil. Int J Syst Evol Microbiol, 69, 3443–3447.

Hyatt, D., Chen, G. L., Locascio, P. F., Land, M. L., Larimer, F. W. & Hauser, L. J. 2010. Prodigal: prokaryotic gene recognition and translation initiation site identification. BMC Bioinformatics, 11, 119.

Ikenaga, M., Kataoka, M., Yin, X., Murouchi, A. & Sakai, M. 2021. Characterization and Distribution of Agar-degrading Steroidobacter agaridevorans sp. nov., Isolated from Rhizosphere Soils. Microbes Environ, 36.

Jain, C., Rodriguez, R. L., Phillippy, A. M., Konstantinidis, K. T. & Aluru, S. 2018. High throughput ANI analysis of 90K prokaryotic genomes reveals clear species boundaries. Nat Commun, 9, 5114.

Kalam, S., Basu, A., Ahmad, I., Sayyed, R. Z., El-Enshasy, H. A., Dailin, D. J. & Suriani, N. L. 2020. Recent Understanding of Soil Acidobacteria and Their Ecological Significance: A Critical Review. Front Microbiol, 11, 580024.

Kanehisa, M. 2019. Toward understanding the origin and evolution of cellular organisms. Protein Sci, 28, 1947–1951.

Kanehisa, M., Furumichi, M., Sato, Y., Kawashima, M. & Ishiguro-Watanabe, M. 2023. KEGG for taxonomy-based analysis of pathways and genomes. Nucleic Acids Res, 51, D587–D592.

Kanehisa, M. & Goto, S. 2000. KEGG: kyoto encyclopedia of genes and genomes. Nucleic Acids Res, 28, 27–30.

Kang, D. D., Froula, J., Egan, R. & Wang, Z. 2015. Metabat, an efficient tool for accurately reconstructing single genomes from complex microbial communities. Peerj, 3, e1165.

Kang, S. M., Khan, A. L., Hussain, J., Ali, L., Kamran, M., Waqas, M. & Lee, I. J. 2012. Rhizonin A from Burkholderia sp. KCTC11096 and its growth promoting role in lettuce seed germination. Molecules, 17, 7980–8.

Kaur, J. & Sharma, J. 2021. Orchid Root Associated Bacteria: Linchpins or Accessories? Front Plant Sci, 12, 661966.

Kim, D. U., Lee, H., Kim, H., Kim, S. G. & Ka, J. O. 2016. Dongia soli sp. nov., isolated from soil from Dokdo, Korea. Antonie Van Leeuwenhoek, 109, 1397–402.

Kull, T. & Hutchings, M. J. 2006. A comparative analysis of decline in the distribution ranges of orchid species in Estonia and the United Kingdom. Biological Conservation, 129, 31–39.

Langmead, B. & Salzberg, S. L. 2012. Fast gapped-read alignment with Bowtie 2. Nat Methods, 9, 357–9.

Li, D., Liu, C. M., Luo, R., Sadakane, K. & Lam, T. W. 2015. MEGAHIT: an ultra-fast single-node solution for large and complex metagenomics assembly via succinct de Bruijn graph. Bioinformatics, 31, 1674–6.

Li, D., Luo, R., Liu, C. M., Leung, C. M., Ting, H. F., Sadakane, K., Yamashita, H. & Lam, T. W. 2016. MEGAHIT v1.0: A fast and scalable metagenome assembler driven by advanced methodologies and community practices. Methods, 102, 3–11.

Li, H., Handsaker, B., Wysoker, A., Fennell, T., Ruan, J., Homer, N., Marth, G., Abecasis, G. & Durbin, R. 2009. The Sequence Alignment/Map format and SAMtools. Bioinformatics, 25, 2078–2079.

Ling, N., Wang, T. & Kuzyakov, Y. 2022. Rhizosphere bacteriome structure and functions. Nat Commun, 13, 836.

Liu, F., Hewezi, T., Lebeis, S. L., Pantalone, V., Grewal, P. S. & Staton, M. E. 2019. Soil indigenous microbiome and plant genotypes cooperatively modify soybean rhizosphere microbiome assembly. BMC Microbiol, 19, 201.

Liu, Y., Jin, J. H., Liu, Y. H., Zhou, Y. G. & Liu, Z. P. 2010. Dongia mobilis gen. nov., sp. nov., a new member of the family Rhodospirillaceae isolated from a sequencing batch reactor for treatment of malachite green effluent. Int J Syst Evol Microbiol, 60, 2780–2785.

Liu, Y., Xu, Z., Chen, L., Xun, W., Shu, X., Chen, Y., Sun, X., Wang, Z., Ren, Y., Shen, Q. & Zhang, R. 2024. Root colonization by beneficial rhizobacteria. FEMS Microbiol Rev, 48.

Martin, F. M., Uroz, S. & Barker, D. G. 2017. Ancestral alliances: Plant mutualistic symbioses with fungi and bacteria. Science, 356.

Mendes, R., Garbeva, P. & Raaijmakers, J. M. 2013. The rhizosphere microbiome: significance of plant beneficial, plant pathogenic, and human pathogenic microorganisms. FEMS Microbiol Rev, 37, 634–63.

Mikheenko, A., Prjibelski, A., Saveliev, V., Antipov, D. & Gurevich, A. 2018. Versatile genome assembly evaluation with QUAST-LG. Bioinformatics, 34, i142–i150.

Miller, I. J., Rees, E. R., Ross, J., Miller, I., Baxa, J., Lopera, J., Kerby, R. L., Rey, F. E. & Kwan, J. C. 2019. Autometa: automated extraction of microbial genomes from individual shotgun metagenomes. Nucleic Acids Res, 47, e57.

Najera, F., Dippold, M. A., Boy, J., Seguel, O., Koester, M., Stock, S., Merino, C., Kuzyakov, Y. & Matus, F. 2020. Effects of drying/rewetting on soil aggregate dynamics and implications for organic matter turnover. Biology and Fertility of Soils, 56, 893–905.

Narayanan, Z. & Glick, B. R. 2022. Secondary Metabolites Produced by Plant Growth-Promoting Bacterial Endophytes. Microorganisms, 10.

Nurk, S., Meleshko, D., Korobeynikov, A. & Pevzner, P. A. 2017. metaSPAdes: a new versatile metagenomic assembler. Genome Res, 27, 824–834.

O’leary, N. A., Wright, M. W., Brister, J. R., Ciufo, S., Haddad, D., Mcveigh, R., Rajput, B., Robbertse, B., Smith-White, B., Ako-Adjei, D., Astashyn, A., Badretdin, A., Bao, Y., Blinkova, O., Brover, V., Chetvernin, V., Choi, J., Cox, E., Ermolaeva, O., Farrell, C. M., Goldfarb, T., Gupta, T., Haft, D., Hatcher, E., Hlavina, W., Joardar, V. S., Kodali, V. K., Li, W., Maglott, D., Masterson, P., Mcgarvey, K. M., Murphy, M. R., O’neill, K., Pujar, S., Rangwala, S. H., Rausch, D., Riddick, L. D., Schoch, C., Shkeda, A., Storz, S. S., Sun, H., Thibaud-Nissen, F., Tolstoy, I., Tully, R. E., Vatsan, A. R., Wallin, C., Webb, D., Wu, W., Landrum, M. J., Kimchi, A., Tatusova, T., Dicuccio, M., Kitts, P., Murphy, T. D. & Pruitt, K. D. 2016. Reference sequence (RefSeq) database at NCBI: current status, taxonomic expansion, and functional annotation. Nucleic Acids Res, 44, D733–45.

Olanrewaju, O. S., Glick, B. R. & Babalola, O. O. 2017. Mechanisms of action of plant growth promoting bacteria. World J Microbiol Biotechnol, 33, 197.

Palla, M., Turrini, A., Cristani, C., Bonora, L., Pellegrini, D., Primicerio, J., Grassi, A., Hilaj, F., Giovannetti, M. & Agnolucci, M. 2022. Impact of sheep wool residues as soil amendments on olive beneficial symbionts and bacterial diversity. Bioresour Bioprocess, 9, 45.

Parks, D. H., Imelfort, M., Skennerton, C. T., Hugenholtz, P. & Tyson, G. W. 2015. CheckM: assessing the quality of microbial genomes recovered from isolates, single cells, and metagenomes. Genome Res, 25, 1043–55.

Paun, O., Bateman, R. M., Fay, M. F., Hedren, M., Civeyrel, L. & Chase, M. W. 2010. Stable epigenetic effects impact adaptation in allopolyploid orchids (Dactylorhiza: Orchidaceae). Mol Biol Evol, 27, 2465–73.

Paun, O., Bateman, R. M., Fay, M. F., Luna, J. A., Moat, J., Hedren, M. & Chase, M. W. 2011. Altered gene expression and ecological divergence in sibling allopolyploids of Dactylorhiza (Orchidaceae). BMC Evol Biol, 11, 113.

Paun, O., Fay, M. F., Soltis, D. E. & Chase, M. W. 2007. Genetic and epigenetic alterations after hybridization and genome doubling. Taxon, 56, 649–56.

Pieterse, C. M., Zamioudis, C., Berendsen, R. L., Weller, D. M., Van Wees, S. C. & Bakker, P. A. 2014. Induced systemic resistance by beneficial microbes. Annu Rev Phytopathol, 52, 347–75.

Pillon, Y., Fay, M. F., Hedren, M., Bateman, R. M., Devey, D. S., Shipunov, A. B., Van Der Bank, M. & Chase, M. W. 2007. Evolution and temporal diversification of western European polyploid species complexes in Dactylorhiza (Orchidaceae). Taxon, 56, 1185–1208.

Qu, Q., Zhang, Z., Peijnenburg, W., Liu, W., Lu, T., Hu, B., Chen, J., Chen, J., Lin, Z. & Qian, H. 2020. Rhizosphere Microbiome Assembly and Its Impact on Plant Growth. J Agric Food Chem, 68, 5024–5038.

Quiroz-Carreno, S., Munoz-Nunez, E., Silva, F. L., Devotto-Moreno, L., Seigler, D. S., Pastene-Navarrete, E., Cespedes-Acuna, C. L. & Alarcon-Enos, J. 2022. Cyclopeptide Alkaloids from Discaria chacaye (Rhamnaceae) as Result of Symbiosis with Frankia (Actinomycetales). Chem Biodivers, 19, e202200630.

Richardson, A. E. & Simpson, R. J. 2011. Soil microorganisms mediating phosphorus availability update on microbial phosphorus. Plant Physiol, 156, 989–96.

Santoyo, G., Moreno-Hagelsieb, G., Orozco-Mosqueda Mdel, C. & Glick, B. R. 2016. Plant growth-promoting bacterial endophytes. Microbiol Res, 183, 92–9.

Siefert, A., Zillig, K. W., Friesen, M. L. & Strauss, S. Y. 2019. Mutualists Stabilize the Coexistence of Congeneric Legumes. Am Nat, 193, 200–212.

Song, D. D., Ren, L., Li, X., Ma, D. L. & Zang, S. Y. 2021. Soil Bacterial Diversity and Composition of Different Forest Types in Greater Xing’an Mountains, China. Applied Ecology and Environmental Research, 19, 1983–1997.

Soó, R. D. 1980. Dactylorhiza. *In*: Tutin, T. G., Heywood, V. H., Burges, N. A., Moore, D. M., Valentine, D. H., Walters, S. M. & Webb, D. A. (eds.) Flora Europea. Cambridge, UK: Cambridge University Press

Timilsena, P. R., Barrett, C. F., Pineyro-Nelson, A., Wafula, E. K., Ayyampalayam, S., Mcneal, J. R., Yukawa, T., Givnish, T. J., Graham, S. W., Pires, J. C., Davis, J. I., Ane, C., Stevenson, D. W., Leebens-Mack, J., Martinez-Salas, E., Alvarez-Buylla, E. R. & Depamphilis, C. W. 2023. Phylotranscriptomic Analyses of Mycoheterotrophic Monocots Show a Continuum of Convergent Evolutionary Changes in Expressed Nuclear Genes From Three Independent Nonphotosynthetic Lineages. Genome Biol Evol, 15.

Toronen, P., Medlar, A. & Holm, L. 2018. PANNZER2: a rapid functional annotation web server. Nucleic Acids Res, 46, W84–W88.

Trivedi, P., Leach, J. E., Tringe, S. G., Sa, T. & Singh, B. K. 2020. Plant-microbiome interactions: from community assembly to plant health. Nat Rev Microbiol, 18, 607–621.

Trivedi, P., Trivedi, C., Grinyer, J., Anderson, I. C. & Singh, B. K. 2016. Harnessing Host-Vector Microbiome for Sustainable Plant Disease Management of Phloem-Limited Bacteria. Front Plant Sci, 7, 1423.

Uroz, S., Courty, P. E. & Oger, P. 2019. Plant Symbionts Are Engineers of the Plant-Associated Microbiome. Trends Plant Sci, 24, 905–916.

Vandenkoornhuyse, P., Quaiser, A., Duhamel, M., Le Van, A. & Dufresne, A. 2015. The importance of the microbiome of the plant holobiont. New Phytol, 206, 1196–206.

Vives-Peris, V., De Ollas, C., Gomez-Cadenas, A. & Perez-Clemente, R. M. 2020. Root exudates: from plant to rhizosphere and beyond. Plant Cell Rep, 39, 3–17.

Walia, A., Mehta, P., Chauhan, A. & Shirkot, C. K. 2013. Effect of Bacillus subtilis Strain CKT1 as Inoculum on Growth of Tomato Seedlings Under Net House Conditions. Proceedings of the National Academy of Sciences, India Section B: Biological Sciences, 84, 145–155.

Walker, T. S., Bais, H. P., Grotewold, E. & Vivanco, J. M. 2003. Root exudation and rhizosphere biology. Plant Physiol, 132, 44–51.

Ward, N. L., Challacombe, J. F., Janssen, P. H., Henrissat, B., Coutinho, P. M., Wu, M., Xie, G., Haft, D. H., Sait, M., Badger, J., Barabote, R. D., Bradley, B., Brettin, T. S., Brinkac, L. M., Bruce, D., Creasy, T., Daugherty, S. C., Davidsen, T. M., Deboy, R. T., Detter, J. C., Dodson, R. J., Durkin, A. S., Ganapathy, A., Gwinn-Giglio, M., Han, C. S., Khouri, H., Kiss, H., Kothari, S. P., Madupu, R., Nelson, K. E., Nelson, W. C., Paulsen, I., Penn, K., Ren, Q., Rosovitz, M. J., Selengut, J. D., Shrivastava, S., Sullivan, S. A., Tapia, R., Thompson, L. S., Watkins, K. L., Yang, Q., Yu, C., Zafar, N., Zhou, L. & Kuske, C. R. 2009. Three genomes from the phylum Acidobacteria provide insight into the lifestyles of these microorganisms in soils. Appl Environ Microbiol, 75, 2046–56.

Wolfe, T. M., Balao, F., Trucchi, E., Bachmann, G., Gu, W., Baar, J., Hedren, M., Weckwerth, W., Leitch, A. R. & Paun, O. 2023. Recurrent allopolyploidizations diversify ecophysiological traits in marsh orchids (Dactylorhiza majalis s.l.). Mol Ecol, 32, 4777–4790.

Yu, G. 2020. Using ggtree to Visualize Data on Tree-Like Structures. Curr Protoc Bioinformatics, 69, e96.

Yu, G., Lam, T. T., Zhu, H. & Guan, Y. 2018. Two Methods for Mapping and Visualizing Associated Data on Phylogeny Using Ggtree. Mol Biol Evol, 35, 3041–3043.

Yu, G., Smith, D. K., Zhu, H., Guan, Y., Lam, T. T. Y. & Mcinerny, G. 2016. GGTREE:an R package for visualization and annotation of phylogenetic trees with their covariates and other associated data. Methods in Ecology and Evolution, 8, 28–36.

## REFERENCES

Albertsen, M., Hugenholtz, P., Skarshewski, A., Nielsen, K. L., Tyson, G. W. & Nielsen, P. H. 2013. Genome sequences of rare, uncultured bacteria obtained by differential coverage binning of multiple metagenomes. Nat Biotechnol, 31, 533–8.

Altshuler, I., Hamel, J., Turney, S., Magnuson, E., Levesque, R., Greer, C. W. & Whyte, L. G. 2019. Species interactions and distinct microbial communities in high Arctic permafrost affected cryosols are associated with the CH(4) and CO(2) gas fluxes. Environ Microbiol, 21, 3711–3727.

Anderson, I., Sikorski, J., Zeytun, A., Nolan, M., Lapidus, A., Lucas, S., Hammon, N., Deshpande, S., Cheng, J. F., Tapia, R., Han, C., Goodwin, L., Pitluck, S., Liolios, K., Pagani, I., Ivanova, N., Huntemann, M., Mavromatis, K., Ovchinikova, G., Pati, A., Chen, A., Palaniappan, K., Land, M., Hauser, L., Brambilla, E. M., Ngatchhou-Djao, O. D., Rohde, M., Tindall, B. J., Goker, M., Detter, J. C., Woyke, T., Bristow, J., Eisen, J. A., Markowitz, V., Hugenholtz, P., Klenk, H. P. & Kyrpides, N. C. 2011. Complete genome sequence of Nitratifractor salsuginis type strain (E9I37-1). Stand Genomic Sci, 4, 322–30.

Asem, M. D., Shi, L., Jiao, J. Y., Wang, D., Han, M. X., Dong, L., Liu, F., Salam, N. & Li, W. J. 2018. Desertimonas flava gen. nov., sp. nov. isolated from a desert soil, and proposal of Ilumatobacteraceae fam. nov. Int J Syst Evol Microbiol, 68, 3593–3599.

Audic, S., Lescot, M., Claverie, J. M. & Scholz, H. C. 2009. Brucella microti: the genome sequence of an emerging pathogen. BMC Genomics, 10, 352.

Badger, J. H., Hoover, T. R., Brun, Y. V., Weiner, R. M., Laub, M. T., Alexandre, G., Mrazek, J., Ren, Q., Paulsen, I. T., Nelson, K. E., Khouri, H. M., Radune, D., Sosa, J., Dodson, R. J., Sullivan, S. A., Rosovitz, M. J., Madupu, R., Brinkac, L. M., Durkin, A. S., Daugherty, S. C., Kothari, S. P., Giglio, M. G., Zhou, L., Haft D. H., Selengut, J. D., Davidsen, T. M., Yang, Q., Zafar, N. & Ward, N. L. 2006. Comparative genomic evidence for a close relationship between the dimorphic prosthecate bacteria Hyphomonas neptunium and Caulobacter crescentus. J Bacteriol, 188, 6841–50.

Baek, C., Shin, S. K. & Yi, H. 2019. Phenylobacterium parvum sp. nov., isolated from lake water. Int J Syst Evol Microbiol, 69, 1169–1172.

Bai, Y., Muller, D. B., Srinivas, G., Garrido-Oter, R., Potthoff, E., Rott, M., Dombrowski, N., Munch, P. C., Spaepen, S., Remus-Emsermann, M., Huttel, B., Mchardy, A. C., Vorholt, J. A. & Schulze-Lefert, P. 2015. Functional overlap of the Arabidopsis leaf and root microbiota. Nature, 528, 364–9.

Behrendt, U., Schumann, P., Hamada, M., Suzuki, K. I., Sproer, C. & Ulrich, A. 2011. Reclassification of Leifsonia ginsengi (Qiu et al. 2007) as Herbiconiux ginsengi gen. nov., comb. nov. and description of Herbiconiux solani sp. nov., an actinobacterium associated with the phyllosphere of Solanum tuberosum L. Int J Syst Evol Microbiol, 61, 1039–1047.

Benaud, N., Edwards, R. J., Amos, T. G., D’agostino, P. M., Gutierrez-Chavez, C., Montgomery, K., Nicetic, I. & Ferrari, B. C. 2021. Antarctic desert soil bacteria exhibit high novel natural product potential, evaluated through long-read genome sequencing and comparative genomics. Environ Microbiol, 23, 3646–3664.

Blattner, F. R., Plunkett, G., 3rd, Bloch, C. A., Perna, N. T., Burland, V., Riley, M., Collado-Vides, J., Glasner, J. D., Rode, C. K., Mayhew, G. F., Gregor, J., Davis, N. W., Kirkpatrick, H. A., Goeden, M. A., Rose, D. J., Mau, B. & Shao, Y. 1997. The complete genome sequence of Escherichia coli K-12. Science, 277, 1453–62.

Boeuf, D., Edwards, B. R., Eppley, J. M., Hu, S. K., Poff, K. E., Romano, A. E., Caron, D. A., Karl, D. M. & Delong, E. F. 2019. Biological composition and microbial dynamics of sinking particulate organic matter at abyssal depths in the oligotrophic open ocean. Proc Natl Acad Sci U S A, 116, 11824–11832.

Bondoso, J., Albuquerque, L., Nobre, M. F., Lobo-Da-Cunha, A., Da Costa, M. S. & Lage, O. M. 2011. Aquisphaera giovannonii gen. nov., sp. nov., a planctomycete isolated from a freshwater aquarium. Int J Syst Evol Microbiol, 61, 2844–2850.

Brown, A. M., Howe, D. K., Wasala, S. K., Peetz, A. B., Zasada, I. A. & Denver, D. R. 2015. Comparative Genomics of a Plant-Parasitic Nematode Endosymbiont Suggest a Role in Nutritional Symbiosis. Genome Biol Evol, 7, 2727–46.

Cabello-Yeves, P. J., Ghai, R., Mehrshad, M., Picazo, A., Camacho, A. & Rodriguez-Valera, F. 2017. Reconstruction of Diverse Verrucomicrobial Genomes from Metagenome Datasets of Freshwater Reservoirs. Front Microbiol, 8, 2131.

Cabello-Yeves, P. J., Zemskaya, T. I., Rosselli, R., Coutinho, F. H., Zakharenko, A. S., Blinov, V. V. & Rodriguez-Valera, F. 2018. Genomes of Novel Microbial Lineages Assembled from the Sub-Ice Waters of Lake Baikal. Appl Environ Microbiol, 84.

Cai, H., Shi, Y., Wang, Y., Cui, H. & Jiang, H. 2017. Aquidulcibacter paucihalophilus gen. nov., sp. nov., a novel member of family Caulobacteraceae isolated from cyanobacterial aggregates in a eutrophic lake. Antonie Van Leeuwenhoek, 110, 1169–1177.

Chen, W. M., Chen, T. Y., Yang, C. C. & Sheu, S. Y. 2020. Oleiharenicola lentus sp. nov., isolated from irrigation water. Int J Syst Evol Microbiol, 70, 3440–3448.

Choi, G. M., Liu, Q., Liu, Q., Jun, M. O., Choi, W. J., Yong Kim, S., Wee, J. H. & Im, W. T. 2022. Hanamia caeni gen. nov., sp. nov., a Member of the Family Chitinophagaceae Isolated from Activated Sludge in Korea. Curr Microbiol, 79, 134.

Christopherson, M. R., Suen, G., Bramhacharya, S., Jewell, K. A., Aylward, F. O., Mead, D. & Brumm, P. J. 2013. The genome sequences of Cellulomonas fimi and “Cellvibrio gilvus” reveal the cellulolytic strategies of two facultative anaerobes, transfer of “Cellvibrio gilvus” to the genus Cellulomonas, and proposal of Cellulomonas gilvus sp. nov. PLoS One, 8, e53954.

Cole, S. T., Brosch, R., Parkhill, J., Garnier, T., Churcher, C., Harris, D., Gordon, S. V., Eiglmeier, K., Gas, S., Barry, C. E., 3rd, Tekaia, F., Badcock, K., Basham, D., Brown, D., Chillingworth, T., Connor, R., Davies, R., Devlin, K., Feltwell, T., Gentles, S., Hamlin, N., Holroyd, S., Hornsby, T., Jagels, K., Krogh, A., Mclean, J., Moule, S., Murphy, L., Oliver, K., Osborne, J., Quail, M. A., Rajandream, M. A., Rogers, J., Rutter, S., Seeger, K., Skelton, J., Squares, R., Squares, S., Sulston, J. E., Taylor, K., Whitehead, S. & Barrell, B. G. 1998. Deciphering the biology of Mycobacterium tuberculosis from the complete genome sequence. Nature, 393, 537–44.

Cole, S. T., Eiglmeier, K., Parkhill, J., James, K. D., Thomson, N. R., Wheeler, P. R., Honore, N., Garnier, T., Churcher, C., Harris, D., Mungall, K., Basham, D., Brown, D., Chillingworth, T., Connor, R., Davies, R. M., Devlin, K., Duthoy, S., Feltwell, T., Fraser, A., Hamlin, N., Holroyd, S., Hornsby, T., Jagels, K., Lacroix, C., Maclean, J., Moule, S., Murphy, L., Oliver, K., Quail, M. A., Rajandream, M. A., Rutherford, K. M., Rutter, S., Seeger, K., Simon, S., Simmonds, M., Skelton, J., Squares, R., Squares, S., Stevens, K., Taylor, K., Whitehead, S., Woodward, J. R. & Barrell, B. G. 2001. Massive gene decay in the leprosy bacillus. Nature, 409, 1007–11.

Collingro, A., Tischler, P., Weinmaier, T., Penz, T., Heinz, E., Brunham, R. C., Read, T. D., Bavoil, P. M., Sachse, K., Kahane, S., Friedman, M. G., Rattei, T., Myers, G. S. & Horn, M. 2011. Unity in variety--the pan-genome of the Chlamydiae. Mol Biol Evol, 28, 3253–70.

Conners, E. M., Davenport, E. J. & Bose, A. 2021. Revised Draft Genome Sequences of Rhodomicrobium vannielii ATCC 17100 and Rhodomicrobium udaipurense JA643. Microbiol Resour Announc, 10.

Dang, X., Xie, Z., Liu, W., Sun, Y., Liu, X., Zhu, Y. & Staehelin, C. 2019. The genome of Ensifer alkalisoli YIC4027 provides insights for host specificity and environmental adaptations. BMC Genomics, 20, 643.

Debruyn, J. M., Radosevich, M., Wommack, K. E., Polson, S. W., Hauser, L. J., Fawaz, M. N., Korlach, J. & Tsai, Y. C. 2014. Genome Sequence and Methylome of Soil Bacterium Gemmatirosa kalamazoonensis KBS708t, a Member of the Rarely Cultivated Gemmatimonadetes Phylum. Genome Announc, 2.

Dedysh, S. N., Kulichevskaya, I. S., Beletsky, A. V., Ivanova, A. A., Rijpstra, W. I. C., Damste, J. S. S., Mardanov, A. V. & Ravin, N. V. 2020. Lacipirellula parvula gen. nov., sp. nov., representing a lineage of planctomycetes widespread in low-oxygen habitats, description of the family Lacipirellulaceae fam. nov. and proposal of the orders Pirellulales ord. nov., Gemmatales ord. nov. and Isosphaerales ord. nov. Syst Appl Microbiol, 43, 126050.

Doroghazi, J. R., Albright, J. C., Goering, A. W., Ju, K. S., Haines, R. R., Tchalukov, K. A., Labeda, D. P., Kelleher, N. L. & Metcalf, W. W. 2014. A roadmap for natural product discovery based on large-scale genomics and metabolomics. Nat Chem Biol, 10, 963–8.

Elcheninov, A. G., Menzel, P., Gudbergsdottir, S. R., Slesarev, A. I., Kadnikov, V. V., Krogh, A., Bonch-Osmolovskaya, E. A., Peng, X. & Kublanov, I. V. 2017. Sugar Metabolism of the First Thermophilic Planctomycete Thermogutta terrifontis: Comparative Genomic and Transcriptomic Approaches. Front Microbiol, 8, 2140.

Eustaquio, A. S., Mcglinchey, R. P., Liu, Y., Hazzard, C., Beer, L. L., Florova, G., Alhamadsheh, M. M., Lechner, A., Kale, A. J., Kobayashi, Y., Reynolds, K. A. & Moore, B. S. 2009. Biosynthesis of the salinosporamide A polyketide synthase substrate chloroethylmalonyl-coenzyme A from S-adenosyl-L-methionine. Proc Natl Acad Sci U S A, 106, 12295–300.

Evtushenko, L. I., Dorofeeva, L. V., Dobrovolskaya, T. G., Streshinskaya, G. M., Subbotin, S. A. & Tiedje, J. M. 2001. Agreia bicolorata gen. nov., sp. nov., to accommodate actinobacteria isolated from narrow reed grass infected by the nematode Heteroanguina graminophila. Int J Syst Evol Microbiol, 51, 2073–2079.

Gao, R. & Sun, C. 2021. A marine bacterial community capable of degrading poly(ethylene terephthalate) and polyethylene. J Hazard Mater, 416, 125928.

Golyshina, O. V., Lunsdorf, H., Kublanov, I. V., Goldenstein, N. I., Hinrichs, K. U. & Golyshin, P. N. 2016. The novel extremely acidophilic, cell-wall-deficient archaeon Cuniculiplasma divulgatum gen. nov., sp. nov. represents a new family, Cuniculiplasmataceae fam. nov., of the order Thermoplasmatales. Int J Syst Evol Microbiol, 66, 332–340.

Gonzalez, D., Huber, K. J., Tindall, B., Hedrich, S., Rojas-Villalobos, C., Quatrini, R., Dinamarca, M. A., Ibacache-Quiroga, C., Schwarz, A., Canales, C. & Nancucheo, I. 2020. Acidiferrimicrobium australe gen. nov., sp. nov., an acidophilic and obligately heterotrophic, member of the Actinobacteria that catalyses dissimilatory oxido-reduction of iron isolated from metal-rich acidic water in Chile. Int J Syst Evol Microbiol, 70, 3348–3354.

Guo, M., Han, X., Jin, T., Zhou, L., Yang, J., Li, Z., Chen, J., Geng, B., Zou, Y., Wan, D., Li, D., Dai, W., Wang, H., Chen, Y., Ni, P., Fang, C. & Yang, R. 2012. Genome sequences of three species in the family Planctomycetaceae. J Bacteriol, 194, 3740–1.

Haack, F. S., Poehlein, A., Kroger, C., Voigt, C. A., Piepenbring, M., Bode, H. B., Daniel, R., Schafer, W. & Streit, W. R. 2016. Molecular Keys to the Janthinobacterium and Duganella spp. Interaction with the Plant Pathogen Fusarium graminearum. Front Microbiol, 7, 1668.

Hassan, Y. I., Lepp, D., He, J. & Zhou, T. 2014. Draft Genome Sequences of Devosia sp. Strain 17-2-E-8 and Devosia riboflavina Strain IFO13584. Genome Announc, 2.

He, C., Keren, R., Whittaker, M., Farag, I. F., Doudna, J., Cate, J. H. D. & Banfield, J. 2020a. Huge and variable diversity of episymbiotic CPR bacteria and DPANN archaea in groundwater ecosystems. bioRxiv, 2020.05.14.094862.

He, Y. Q., Chen, R. W., Li, C., Shi, S. B., Cui, L. Q., Long, L. J. & Tian, X. P. 2020b. Actinomarinicola tropica gen. nov. sp. nov., a new marine actinobacterium of the family Iamiaceae, isolated from South China Sea sediment environments. Int J Syst Evol Microbiol, 70, 3852–3858.

Higashioka, Y., Kojima, H., Watanabe, M. & Fukui, M. 2013. Desulfatitalea tepidiphila gen. nov., sp. nov., a sulfate-reducing bacterium isolated from tidal flat sediment. Int J Syst Evol Microbiol, 63, 761–765.

Hu, Z. Y., Wang, Y. Z., Im, W. T., Wang, S. Y., Zhao, G. P., Zheng, H. J. & Quan, Z. X. 2014. The first complete genome sequence of the class Fimbriimonadia in the phylum Armatimonadetes. PLoS One, 9, e100794.

Ivanova, N., Daum, C., Lang, E., Abt, B., Kopitz, M., Saunders, E., Lapidus, A., Lucas, S., Glavina Del Rio, T., Nolan, M., Tice, H., Copeland, A., Cheng, J. F., Chen, F., Bruce, D., Goodwin, L., Pitluck, S., Mavromatis, K., Pati, A., Mikhailova, N., Chen, A., Palaniappan, K., Land, M., Hauser, L., Chang, Y. J., Jeffries, C. D., Detter, J. C., Brettin, T., Rohde, M., Goker, M., Bristow, J., Markowitz, V., Eisen, J. A., Hugenholtz, P., Kyrpides, N. C. & Klenk, H. P. 2010. Complete genome sequence of Haliangium ochraceum type strain (SMP-2). Stand Genomic Sci, 2, 96–106.

Ivanova, N., Sorokin, A., Anderson, I., Galleron, N., Candelon, B., Kapatral, V., Bhattacharyya, A., Reznik, G., Mikhailova, N., Lapidus, A., Chu, L., Mazur, M., Goltsman, E., Larsen, N., D’souza, M., Walunas, T., Grechkin, Y., Pusch, G., Haselkorn, R., Fonstein, M., Ehrlich, S. D., Overbeek, R. & Kyrpides, N. 2003. Genome sequence of Bacillus cereus and comparative analysis with Bacillus anthracis. Nature, 423, 87–91.

Ji, M., Greening, C., Vanwonterghem, I., Carere, C. R., Bay, S. K., Steen, J. A., Montgomery, K., Lines, T., Beardall, J., Van Dorst, J., Snape, I., Stott, M. B., Hugenholtz, P. & Ferrari, B. C. 2017. Atmospheric trace gases support primary production in Antarctic desert surface soil. Nature, 552, 400–403.

Kallscheuer, N., Rast, P., Jogler, M., Wiegand, S., Kohn, T., Boedeker, C., Jeske, O., Heuer, A., Quast, C., Glockner, F. O., Rohde, M. & Jogler, C. 2021. Analysis of bacterial communities in a municipal duck pond during a phytoplankton bloom and isolation of Anatilimnocola aggregata gen. nov., sp. nov., Lacipirellula limnantheis sp. nov. and Urbifossiella limnaea gen. nov., sp. nov. belonging to the phylum Planctomycetes. Environ Microbiol, 23, 1379–1396.

Kallscheuer, N., Wiegand, S., Heuer, A., Rensink, S., Boersma, A. S., Jogler, M., Boedeker, C., Peeters, S. H., Rast, P., Jetten, M. S. M., Rohde, M. & Jogler, C. 2020. Blastopirellula retiformator sp. nov. isolated from the shallow-sea hydrothermal vent system close to Panarea Island. Antonie Van Leeuwenhoek, 113, 1811–1822.

Kaneko, T., Maita, H., Hirakawa, H., Uchiike, N., Minamisawa, K., Watanabe, A. & Sato, S. 2011. Complete Genome Sequence of the Soybean Symbiont Bradyrhizobium japonicum Strain USDA6T. Genes (Basel), 2, 763–87.

Kaneko, T., Nakamura, Y., Sato, S., Minamisawa, K., Uchiumi, T., Sasamoto, S., Watanabe, A., Idesawa, K., Iriguchi, M., Kawashima, K., Kohara, M., Matsumoto, M., Shimpo, S., Tsuruoka, H., Wada, T., Yamada, M. & Tabata, S. 2002. Complete genomic sequence of nitrogen-fixing symbiotic bacterium Bradyrhizobium japonicum USDA110. DNA Res, 9, 189–97.

Kantor, R. S., Wrighton, K. C., Handley, K. M., Sharon, I., Hug, L. A., Castelle, C. J., Thomas, B. C. & Banfield, J. F. 2013. Small genomes and sparse metabolisms of sediment-associated bacteria from four candidate phyla. mBio, 4, e00708–13.

Kawaichi, S., Ito, N., Kamikawa, R., Sugawara, T., Yoshida, T. & Sako, Y. 2013. Ardenticatena maritima gen. nov., sp. nov., a ferric iron- and nitrate-reducing bacterium of the phylum ’Chloroflexi’ isolated from an iron-rich coastal hydrothermal field, and description of Ardenticatenia classis nov. Int J Syst Evol Microbiol, 63, 2992–3002.

Khaleque, H. N., Gonzalez, C., Kaksonen, A. H., Boxall, N. J., Holmes, D. S. & Watkin, E. L. J. 2019. Genome-based classification of two halotolerant extreme acidophiles, Acidihalobacter prosperus V6 (=DSM 14174 =JCM 32253) and ’Acidihalobacter ferrooxidans’ V8 (=DSM 14175 =JCM 32254) as two new species, Acidihalobacter aeolianus sp. nov. and Acidihalobacter ferrooxydans sp. nov., respectively. Int J Syst Evol Microbiol, 69, 1557–1565.

Kropp, K. G., Davidova, I. A. & Suflita, J. M. 2000. Anaerobic oxidation of n-dodecane by an addition reaction in a sulfate-reducing bacterial enrichment culture. Appl Environ Microbiol, 66, 5393–8.

Kulichevskaya, I. S., Naumoff, D. G., Miroshnikov, K. K., Ivanova, A. A., Philippov, D. A., Hakobyan, A., Rijpstra, W. I. C., Damste, J. S. S., Liesack, W. & Dedysh, S. N. 2020. Limnoglobus roseus gen. nov., sp. nov., a novel freshwater planctomycete with a giant genome from the family Gemmataceae. Int J Syst Evol Microbiol, 70, 1240–1249.

Kumar, G., Lhingjakim, K. L., Uppada, J., Ahamad, S., Kumar, D., Kashif, G. M., Sasikala, C. & Ramana, C. V. 2021. Aquisphaera insulae sp. nov., a new member in the family Isosphaeraceae, isolated from the floating island of Loktak lake and emended description of the genus Aquisphaera. Antonie Van Leeuwenhoek, 114, 1465–1477.

Kunst, F., Ogasawara, N., Moszer, I., Albertini, A. M., Alloni, G., Azevedo, V., Bertero, M. G., Bessieres, P., Bolotin, A., Borchert, S., Borriss, R., Boursier, L., Brans, A., Braun, M., Brignell, S. C., Bron, S., Brouillet, S., Bruschi, C. V., Caldwell, B., Capuano, V., Carter, N. M., Choi, S. K., Cordani, J. J., Connerton, I. F., Cummings, N. J., Daniel, R. A., Denziot, F., Devine, K. M., Dusterhoft, A., Ehrlich, S. D., Emmerson, P. T., Entian, K. D., Errington, J., Fabret, C., Ferrari, E., Foulger, D., Fritz, C., Fujita, M., Fujita, Y., Fuma, S., Galizzi, A., Galleron, N., Ghim, S. Y., Glaser, P., Goffeau, A., Golightly, E. J., Grandi, G., Guiseppi, G., Guy, B. J., Haga, K., Haiech, J., Harwood, C. R., Henaut, A., Hilbert, H., Holsappel, S., Hosono, S., Hullo, M. F., Itaya, M., Jones, L., Joris, B., Karamata, D., Kasahara, Y., Klaerr-Blanchard, M., Klein, C., Kobayashi, Y., Koetter, P., Koningstein, G., Krogh, S., Kumano, M., Kurita, K., Lapidus, A., Lardinois, S., Lauber, J., Lazarevic, V., Lee, S. M., Levine, A., Liu, H., Masuda, S., Mauel, C., Medigue, C., Medina, N., Mellado, R. P., Mizuno, M., Moestl, D., Nakai, S., Noback, M., Noone, D., O’reilly, M., Ogawa, K., Ogiwara, A., Oudega, B., Park, S. H., Parro, V., Pohl, T. M., Portelle, D., Porwollik, S., Prescott, A. M., Presecan, E., Pujic, P., Purnelle, B., et al. 1997. The complete genome sequence of the gram-positive bacterium Bacillus subtilis. Nature, 390, 249–56.

Kyrpides, N. C., Woyke, T., Eisen, J. A., Garrity, G., Lilburn, T. G., Beck, B. J., Whitman, W. B., Hugenholtz, P. & Klenk, H. P. 2014. Genomic Encyclopedia of Type Strains, Phase I: The one thousand microbial genomes (KMG-I) project. Stand Genomic Sci, 9, 1278–84.

Lasarre, B., Kysela, D. T., Stein, B. D., Ducret, A., Brun, Y. V. & Mckinlay, J. B. 2018. Restricted Localization of Photosynthetic Intracytoplasmic Membranes (ICMs) in Multiple Genera of Purple Nonsulfur Bacteria. mBio, 9.

Lauro, F. M., Mcdougald, D., Thomas, T., Williams, T. J., Egan, S., Rice, S., Demaere, M. Z., Ting, L., Ertan, H., Johnson, J., Ferriera, S., Lapidus, A., Anderson, I., Kyrpides, N., Munk, A. C., Detter, C., Han, C. S., Brown, M. V., Robb, F. T., Kjelleberg, S. & Cavicchioli, R. 2009. The genomic basis of trophic strategy in marine bacteria. Proc Natl Acad Sci U S A, 106, 15527–33.

Lawson, C. E., Wu, S., Bhattacharjee, A. S., Hamilton, J. J., Mcmahon, K. D., Goel, R. & Noguera, D. R. 2017. Metabolic network analysis reveals microbial community interactions in anammox granules. Nat Commun, 8, 15416.

Lee, J. S., Lee, K. C., Kim, K. K. & Lee, B. 2016. Complete genome sequence of the Variibacter gotjawalensis GJW-30(T) from soil of lava forest, Gotjawal. J Biotechnol, 218, 64–5.

Lee, K. C., Morgan, X. C., Dunfield, P. F., Tamas, I., Mcdonald, I. R. & Stott, M. B. 2014. Genomic analysis of Chthonomonas calidirosea, the first sequenced isolate of the phylum Armatimonadetes. ISME J, 8, 1522–33.

Lee, S. A., Kanth, B. K., Kim, H. S., Kim, T. W., Sang, M. K., Song, J. & Weon, H. Y. 2019a. Complete genome sequence of the plant growth-promoting endophytic bacterium Rhodanobacter glycinis T01E-68 isolated from tomato (Solanum lycopersicum L.) plant roots. Korean J. Microbiol., 55, 422–424.

Lee, S. A., Kim, Y., Sang, M. K., Song, J., Kwon, S. W. & Weon, H. Y. 2019b. Chryseolinea soli sp. nov., isolated from soil. J Microbiol, 57, 122–126.

Lim, Y. K., Park, S. N., Lee, W. P., Shin, J. H., Jo, E., Shin, Y., Paek, J., Chang, Y. H., Kim, H. & Kook, J. K. 2019. Lautropia dentalis sp. nov., Isolated from Human Dental Plaque of a Gingivitis Lesion. Curr Microbiol, 76, 1369–1373.

Liu, L., Wang, Y., Che, Y., Chen, Y., Xia, Y., Luo, R., Cheng, S. H., Zheng, C. & Zhang, T. 2020. High-quality bacterial genomes of a partial-nitritation/anammox system by an iterative hybrid assembly method. Microbiome, 8, 155.

Liu, Z. P., Wang, B. J., Liu, S. J. & Liu, Y. H. 2005. Asticcacaulis taihuensis sp. nov., a novel stalked bacterium isolated from Taihu Lake, China. Int J Syst Evol Microbiol, 55, 1239–1242.

Liu, Z. T., Jiao, J. Y., Liu, L., Li, M. M., Ming, Y. Z., Song, J. L., Lv, A. P., Xian, W. D., Fang, B. Z. & Li, W. J. 2019. Rhabdothermincola sediminis gen. nov., sp. nov., a new actinobacterium isolated from hot spring sediment, and emended description of the family Iamiaceae. Int J Syst Evol Microbiol, 71.

Lopez, G., Diaz-Cardenas, C., David Alzate, J., Gonzalez, L. N., Shapiro, N., Woyke, T., Kyrpides, N. C., Restrepo, S. & Baena, S. 2018. Description of Alicyclobacillus montanus sp. nov., a mixotrophic bacterium isolated from acidic hot springs. Int J Syst Evol Microbiol, 68, 1608–1615.

Mathew, M. J., Subramanian, G., Nguyen, T. T., Robert, C., Mediannikov, O., Fournier, P. E. & Raoult, D. 2012. Genome sequence of Diplorickettsia massiliensis, an emerging Ixodes ricinus-associated human pathogen. J Bacteriol, 194, 3287.

Matsuura, N., Tourlousse, D. M., Ohashi, A., Hugenholtz, P. & Sekiguchi, Y. 2015. Draft Genome Sequences of Anaerolinea thermolimosa IMO-1, Bellilinea caldifistulae GOMI-1, Leptolinea tardivitalis YMTK-2, Levilinea saccharolytica KIBI-1, Longilinea arvoryzae KOME-1, Previously Described as Members of the Class Anaerolineae (Chloroflexi). Genome Announc, 3.

Maurais, E. G., Iannazzi, L. C. & Maclea, K. S. 2022. Genome Sequence of Litorilinea aerophila, an Icelandic Intertidal Hot Springs Bacterium. Microbiol Resour Announc, 11, e0120621.

Mavromatis, K., Sikorski, J., Lapidus, A., Glavina Del Rio, T., Copeland, A., Tice, H., Cheng, J. F., Lucas, S., Chen, F., Nolan, M., Bruce, D., Goodwin, L., Pitluck, S., Ivanova, N., Ovchinnikova, G., Pati, A., Chen, A., Palaniappan, K., Land, M., Hauser, L., Chang, Y. J., Jeffries, C. D., Chain, P., Meincke, L., Sims, D., Chertkov, O., Han, C., Brettin, T., Detter, J. C., Wahrenburg, C., Rohde, M., Pukall, R., Goker, M., Bristow, J., Eisen, J. A., Markowitz, V., Hugenholtz, P., Klenk, H. P. & Kyrpides, N. C. 2010. Complete genome sequence of Alicyclobacillus acidocaldarius type strain (104-IA). Stand Genomic Sci, 2, 9–18.

Miller, T. R., Delcher, A. L., Salzberg, S. L., Saunders, E., Detter, J. C. & Halden, R. U. 2010. Genome sequence of the dioxin-mineralizing bacterium Sphingomonas wittichii RW1. J Bacteriol, 192, 6101–2.

Mino, S., Yoneyama, N., Nakagawa, S., Takai, K. & Sawabe, T. 2018. Enrichment and Genomic Characterization of a N(2)O-Reducing Chemolithoautotroph From a Deep-Sea Hydrothermal Vent. Front Bioeng Biotechnol, 6, 184.

Mo, K. L., Huang, H. Q., Wu, Q. J. & Hu, Y. H. 2020. Amaricoccus solimangrovi sp. nov., isolated from mangrove soil. Int J Syst Evol Microbiol, 70, 5389–5393.

Monteiro-Vitorello, C. B., Zerillo, M. M., Van Sluys, M. A., Camargo, L. E. & Kitajima, J. P. 2013. Complete Genome Sequence of Leifsonia xyli subsp. cynodontis Strain DSM46306, a Gram-Positive Bacterial Pathogen of Grasses. Genome Announc, 1.

Nakahara, N., Nobu, M. K., Takaki, Y., Miyazaki, M., Tasumi, E., Sakai, S., Ogawara, M., Yoshida, N., Tamaki, H., Yamanaka, Y., Katayama, A., Yamaguchi, T., Takai, K. & Imachi, H. 2019. Aggregatilinea lenta gen. nov., sp. nov., a slow-growing, facultatively anaerobic bacterium isolated from subseafloor sediment, and proposal of the new order Aggregatilineales ord. nov. within the class Anaerolineae of the phylum Chloroflexi. Int J Syst Evol Microbiol, 69, 1185–1194.

Nierman, W. C., Feldblyum, T. V., Laub, M. T., Paulsen, I. T., Nelson, K. E., Eisen, J. A., Heidelberg, J. F., Alley, M. R., Ohta, N., Maddock, J. R., Potocka, I., Nelson, W. C., Newton, A., Stephens, C., Phadke, N. D., Ely, B., Deboy, R. T., Dodson, R. J., Durkin, A. S., Gwinn, M. L., Haft, D. H., Kolonay, J. F., Smit, J., Craven, M. B., Khouri, H., Shetty, J., Berry, K., Utterback, T., Tran, K., Wolf, A., Vamathevan, J., Ermolaeva, M., White, O., Salzberg, S. L., Venter, J. C., Shapiro, L. & Fraser, C. M. 2001. Complete genome sequence of Caulobacter crescentus. Proc Natl Acad Sci U S A, 98, 4136–41.

Ormeno-Orrillo, E., Rogel, M. A., Zuniga-Davila, D. & Martinez-Romero, E. 2018. Complete Genome Sequence of the Symbiotic Strain Bradyrhizobium icense LMTR 13(T), Isolated from Lima Bean (Phaseolus lunatus) in Peru. Genome Announc, 6.

Pal, S., Sharma, G. & Subramanian, S. 2021. Complete genome sequence and identification of polyunsaturated fatty acid biosynthesis genes of the myxobacterium Minicystis rosea DSM 24000(T). BMC Genomics, 22, 655.

Parks, D. H., Rinke, C., Chuvochina, M., Chaumeil, P. A., Woodcroft, B. J., Evans, P. N., Hugenholtz, P. & Tyson, G. W. 2017. Recovery of nearly 8,000 metagenome-assembled genomes substantially expands the tree of life. Nat Microbiol, 2, 1533–1542.

Plugge, C. M., Henstra, A. M., Worm, P., Swarts, D. C., Paulitsch-Fuchs, A. H., Scholten, J. C., Lykidis, A., Lapidus, A. L., Goltsman, E., Kim, E., Mcdonald, E., Rohlin, L., Crable, B. R., Gunsalus, R. P., Stams, A. J. & Mcinerney, M. J. 2012. Complete genome sequence of Syntrophobacter fumaroxidans strain (MPOB(T)). Stand Genomic Sci, 7, 91–106.

Pukall, R., Lapidus, A., Glavina Del Rio, T., Copeland, A., Tice, H., Cheng, J. F., Lucas, S., Chen, F., Nolan, M., Bruce, D., Goodwin, L., Pitluck, S., Mavromatis, K., Ivanova, N., Ovchinnikova, G., Pati, A., Chen, A., Palaniappan, K., Land, M., Hauser, L., Chang, Y. J., Jeffries, C. D., Chain, P., Meincke, L., Sims, D., Brettin, T., Detter, J. C., Rohde, M., Goker, M., Bristow, J., Eisen, J. A., Markowitz, V., Kyrpides, N. C., Klenk, H. P. & Hugenholtz, P. 2010. Complete genome sequence of Conexibacter woesei type strain (ID131577). Stand Genomic Sci, 2, 212–9.

Quan, X. T., Siddiqi, M. Z., Liu, Q. Z., Lee, S. M. & Im, W. T. 2020. Devosia ginsengisoli sp. nov., isolated from ginseng cultivation soil. Int J Syst Evol Microbiol, 70, 1489–1495.

Ravin, N. V., Rakitin, A. L., Ivanova, A. A., Beletsky, A. V., Kulichevskaya, I. S., Mardanov, A. V. & Dedysh, S. N. 2018. Genome Analysis of Fimbriiglobus ruber SP5(T), a Planctomycete with Confirmed Chitinolytic Capability. Appl Environ Microbiol, 84.

Read, T. D., Peterson, S. N., Tourasse, N., Baillie, L. W., Paulsen, I. T., Nelson, K. E., Tettelin, H., Fouts, D. E., Eisen, J. A., Gill, S. R., Holtzapple, E. K., Okstad, O. A., Helgason, E., Rilstone, J., Wu, M., Kolonay, J. F., Beanan, M. J., Dodson, R. J., Brinkac, L. M., Gwinn, M., Deboy, R. T., Madpu, R., Daugherty, S. C., Durkin, A. S., Haft, D. H., Nelson, W. C., Peterson, J. D., Pop, M., Khouri, H. M., Radune, D., Benton, J. L., Mahamoud, Y., Jiang, L., Hance, I. R., Weidman, J. F., Berry, K. J., Plaut, R. D., Wolf, A. M., Watkins, K. L., Nierman, W. C., Hazen, A., Cline, R., Redmond, C., Thwaite, J. E., White, O., Salzberg, S. L., Thomason, B., Friedlander, A. M., Koehler, T. M., Hanna, P. C., Kolsto, A. B. & Fraser, C. M. 2003. The genome sequence of Bacillus anthracis Ames and comparison to closely related bacteria. Nature, 423, 81–6.

Reysenbach, A. L., St John, E., Meneghin, J., Flores, G. E., Podar, M., Dombrowski, N., Spang, A., L’haridon, S., Humphris, S. E., De Ronde, C. E. J., Caratori Tontini, F., Tivey, M., Stucker, V. K., Stewart, L. C., Diehl, A. & Bach, W. 2020. Complex subsurface hydrothermal fluid mixing at a submarine arc volcano supports distinct and highly diverse microbial communities. Proc Natl Acad Sci U S A, 117, 32627–32638.

Rinke, C., Schwientek, P., Sczyrba, A., Ivanova, N. N., Anderson, I. J., Cheng, J. F., Darling, A., Malfatti, S., Swan, B. K., Gies, E. A., Dodsworth, J. A., Hedlund, B. P., Tsiamis, G., Sievert, S. M., Liu, W. T., Eisen, J. A., Hallam, S. J., Kyrpides, N. C., Stepanauskas, R., Rubin, E. M., Hugenholtz, P. & Woyke, T. 2013. Insights into the phylogeny and coding potential of microbial dark matter. Nature, 499, 431–7.

Safronova, V. I., Sazanova, A. L., Kuznetsova, I. G., Belimov, A. A., Andronov, E. E., Chirak, E. R., Popova, J. P., Verkhozina, A. V., Willems, A. & Tikhonovich, I. A. 2018. Phyllobacterium zundukense sp. nov., a novel species of rhizobia isolated from root nodules of the legume species Oxytropis triphylla (Pall.) Pers. Int J Syst Evol Microbiol, 68, 1644–1651.

Sakai, H. D. & Kurosawa, N. 2019. Complete genome sequence of the Sulfodiicoccus acidiphilus strain HS-1(T), the first crenarchaeon that lacks polB3, isolated from an acidic hot spring in Ohwaku-dani, Hakone, Japan. BMC Res Notes, 12, 444.

Sangal, V., Goodfellow, M., Jones, A. L., Schwalbe, E. C., Blom, J., Hoskisson, P. A. & Sutcliffe, I. C. 2016. Next-generation systematics: An innovative approach to resolve the structure of complex prokaryotic taxa. Sci Rep, 6, 38392.

Schlesner, H., Rensmann, C., Tindall, B. J., Gade, D., Rabus, R., Pfeiffer, S. & Hirsch, P. 2004. Taxonomic heterogeneity within the Planctomycetales as derived by DNA– DNA hybridization, description of Rhodopirellula baltica gen. nov., sp. nov., transfer of Pirellula marina to the genus Blastopirellula gen. nov. as Blastopirellula marina comb. nov. and emended description of the genus Pirellula. International Journal of Systematic and Evolutionary Microbiology, 54, 1567–1580.

Schneiker, S., Perlova, O., Kaiser, O., Gerth, K., Alici, A., Altmeyer, M. O., Bartels, D., Bekel, T., Beyer, S., Bode, E., Bode, H. B., Bolten, C. J., Choudhuri, J. V., Doss, S., Elnakady, Y. A., Frank, B., Gaigalat, L., Goesmann, A., Groeger, C., Gross, F., Jelsbak, L., Jelsbak, L., Kalinowski, J., Kegler, C., Knauber, T., Konietzny, S., Kopp, M., Krause, L., Krug, D., Linke, B., Mahmud, T., Martinez-Arias, R., Mchardy, A. C., Merai, M., Meyer, F., Mormann, S., Munoz-Dorado, J., Perez, J., Pradella, S., Rachid, S., Raddatz, G., Rosenau, F., Ruckert, C., Sasse, F., Scharfe, M., Schuster, S. C., Suen, G., Treuner-Lange, A., Velicer, G. J., Vorholter, F. J., Weissman, K. J., Welch, R. D., Wenzel, S. C., Whitworth, D. E., Wilhelm, S., Wittmann, C., Blocker, H., Puhler, A. & Muller, R. 2007. Complete genome sequence of the myxobacterium Sorangium cellulosum. Nat Biotechnol, 25, 1281–9.

Seo, J. S., Chong, H., Park, H. S., Yoon, K. O., Jung, C., Kim, J. J., Hong, J. H., Kim, H., Kim, J. H., Kil, J. I., Park, C. J., Oh, H. M., Lee, J. S., Jin, S. J., Um, H. W., Lee, H. J., OH, S. J., Kim, J. Y., Kang, H. L., Lee, S. Y., Lee, K. J. & Kang, H. S. 2005. The genome sequence of the ethanologenic bacterium Zymomonas mobilis ZM4. Nat Biotechnol, 23, 63–8.

Severino, R., Froufe, H. J. C., Barroso, C., Albuquerque, L., Lobo-Da-Cunha, A., Da Costa, M. S. & Egas, C. 2019. High-quality draft genome sequence of Gaiella occulta isolated from a 150 meter deep mineral water borehole and comparison with the genome sequences of other deep-branching lineages of the phylum Actinobacteria. Microbiologyopen, 8, e00840.

Siddiqi, M. Z., Liu, Q., Kang, M. S., Kim, M. S. & Im, W. T. 2016. Anseongella ginsenosidimutans gen. nov., sp. nov., isolated from soil cultivating ginseng. Int J Syst Evol Microbiol, 66, 1125x–1130.

Singleton, C. M., Petriglieri, F., Kristensen, J. M., Kirkegaard, R. H., Michaelsen, T. Y., Andersen, M. H., Kondrotaite, Z., Karst, S. M., Dueholm, M. S., Nielsen, P. H. & Albertsen, M. 2021. Connecting structure to function with the recovery of over 1000 high-quality metagenome-assembled genomes from activated sludge using long-read sequencing. Nat Commun, 12, 2009.

Tamaki, H., Tanaka, Y., Matsuzawa, H., Muramatsu, M., Meng, X. Y., Hanada, S., Mori, K. & Kamagata, Y. 2011. Armatimonas rosea gen. nov., sp. nov., of a novel bacterial phylum, Armatimonadetes phyl. nov., formally called the candidate phylum OP10. Int J Syst Evol Microbiol, 61, 1442–1447.

Tice, H., Mayilraj, S., Sims, D., Lapidus, A., Nolan, M., Lucas, S., Glavina Del Rio, T., Copeland, A., Cheng, J. F., Meincke, L., Bruce, D., Goodwin, L., Pitluck, S., Ivanova, N., Mavromatis, K., Ovchinnikova, G., Pati, A., Chen, A., Palaniappan, K., Land, M., Hauser, L., Chang, Y. J., Jeffries, C. D., Detter, J. C., Brettin, T., Rohde, M., Goker, M., Bristow, J., Eisen, J. A., Markowitz, V., Hugenholtz, P., Kyrpides, N. C., Klenk, H. P. & Chen, F. 2010. Complete genome sequence of Nakamurella multipartita type strain (Y-104). Stand Genomic Sci, 2, 168–75.

Tully, B. J., Sachdeva, R., Graham, E. D. & Heidelberg, J. F. 2017. 290 metagenome-assembled genomes from the Mediterranean Sea: a resource for marine microbiology. Peerj, 5, e3558.

Van Passel, M. W., Kant, R., Palva, A., Copeland, A., Lucas, S., Lapidus, A., Glavina Del Rio, T., Pitluck, S., Goltsman, E., Clum, A., Sun, H., Schmutz, J., Larimer, F. W., Land, M. L., Hauser, L., Kyrpides, N., Mikhailova, N., Richardson, P. P., Janssen, P. H., De Vos, W. M. & Smidt, H. 2011. Genome sequence of the verrucomicrobium Opitutus terrae PB90-1, an abundant inhabitant of rice paddy soil ecosystems. J Bacteriol, 193, 2367–8.

Vieira, S., Pascual, J., Boedeker, C., Geppert, A., Riedel, T., Rohde, M. & Overmann, J. 2020. Terricaulis silvestris gen. nov., sp. nov., a novel prosthecate, budding member of the family Caulobacteraceae isolated from forest soil. Int J Syst Evol Microbiol, 70, 4966–4977.

Wang, J. J., Chen, Q. & Li, Y. Z. 2018. Chryseolinea flava sp. nov., a new species of Chryseolinea isolated from soil. Int J Syst Evol Microbiol, 68, 3518–3522.

Wang, P. H., Chen, Y. L., Wei, S. T., Wu, K., Lee, T. H., Wu, T. Y. & Chiang, Y. R. 2020. Retroconversion of estrogens into androgens by bacteria via a cobalamin-mediated methylation. Proc Natl Acad Sci U S A, 117, 1395–1403.

Welly, B. T., Miller, M. R., Stott, J. L., Blanchard, M. T., Islas-Trejo, A. D., O’rourke, S. M., Young, A. E., Medrano, J. F. & Van Eenennaam, A. L. 2017. Genome Report: Identification and Validation of Antigenic Proteins from Pajaroellobacter abortibovis Using De Novo Genome Sequence Assembly and Reverse Vaccinology. G3 (Bethesda), 7, 321–331.

Whitman, W. B., Woyke, T., Klenk, H. P., Zhou, Y., Lilburn, T. G., Beck, B. J., DE Vos, P., Vandamme, P., Eisen, J. A., Garrity, G., Hugenholtz, P. & Kyrpides, N. C. 2015. Genomic Encyclopedia of Bacterial and Archaeal Type Strains, Phase III: the genomes of soil and plant-associated and newly described type strains. Stand Genomic Sci, 10, 26.

Williams, L. E., Baltrus, D. A., O’donnell, S. D., Skelly, T. J. & Martin, M. O. 2017. Complete Genome Sequence of the Predatory Bacterium Ensifer adhaerens Casida A. Genome Announc, 5.

Woodhouse, J. N., Makower, A. K., Grossart, H. P. & Dittmann, E. 2017. Draft Genome Sequences of Two Uncultured Armatimonadetes Associated with a Microcystis sp. (Cyanobacteria) Isolate. Genome Announc, 5.

Wu, L. & Ma, J. 2019. The Global Catalogue of Microorganisms (GCM) 10K type strain sequencing project: providing services to taxonomists for standard genome sequencing and annotation. Int J Syst Evol Microbiol, 69, 895–898.

Xing, J., Li, X., Sun, Y., Zhao, J., Miao, S., Xiong, Q., Zhang, Y. & Zhang, G. 2019. Comparative genomic and functional analysis of Akkermansia muciniphila and closely related species. Genes Genomics, 41, 1253–1264.

Yamazaki, Y., Akashi, R., Banno, Y., Endo, T., Ezura, H., Fukami-Kobayashi, K., Inaba, K., Isa, T., Kamei, K., Kasai, F., Kobayashi, M., Kurata, N., Kusaba, M., Matuzawa, T., Mitani, S., Nakamura, T., Nakamura, Y., Nakatsuji, N., Naruse, K., Niki, H., Nitasaka, E., Obata, Y., Okamoto, H., Okuma, M., Sato, K., Serikawa, T., Shiroishi, T., Sugawara, H., Urushibara, H., Yamamoto, M., Yaoita, Y., Yoshiki, A. & Kohara, Y. 2010. NBRP databases: databases of biological resources in Japan. Nucleic Acids Res, 38, D26–32.

Yang, F. C., Chen, Y. L., Tang, S. L., Yu, C. P., Wang, P. H., Ismail, W., Wang, C. H., Ding, J. Y., Yang, C. Y., Yang, C. Y. & Chiang, Y. R. 2016. Integrated multi-omics analyses reveal the biochemical mechanisms and phylogenetic relevance of anaerobic androgen biodegradation in the environment. ISME J, 10, 1967–83.

Yee, B., Oertli, G. E., Fuerst, J. A. & Staley, J. T. 2010. Reclassification of the polyphyletic genus Prosthecomicrobium to form two novel genera, Vasilyevaea gen. nov. and Bauldia gen. nov. with four new combinations: Vasilyevaea enhydra comb. nov., Vasilyevaea mishustinii comb. nov., Bauldia consociata comb. nov. and Bauldia litoralis comb. nov. Int J Syst Evol Microbiol, 60, 2960–2966.

Zaburannyi, N., Bunk, B., Maier, J., Overmann, J. & Muller, R. 2016. Genome Analysis of the Fruiting Body-Forming Myxobacterium Chondromyces crocatus Reveals High Potential for Natural Product Biosynthesis. Appl Environ Microbiol, 82, 1945–1957.

Zeng, Y., Baumbach, J., Barbosa, E. G., Azevedo, V., Zhang, C. & Koblizek, M. 2016. Metagenomic evidence for the presence of phototrophic Gemmatimonadetes bacteria in diverse environments. Environ Microbiol Rep, 8, 139–49.

Zhang, H., Sekiguchi, Y., Hanada, S., Hugenholtz, P., Kim, H., Kamagata, Y. & Nakamura, K. 2003. Gemmatimonas aurantiaca gen. nov., sp. nov., a gram-negative, aerobic, polyphosphate-accumulating micro-organism, the first cultured representative of the new bacterial phylum Gemmatimonadetes phyl. nov. Int J Syst Evol Microbiol, 53, 1155–1163.

Zhou, Z., Liu, Y., Xu, W., Pan, J., Luo, Z. H. & Li, M. 2020. Genome- and Community-Level Interaction Insights into Carbon Utilization and Element Cycling Functions of Hydrothermarchaeota in Hydrothermal Sediment. mSystems, 5.

Zhu, L., Long, M., Si, M., Wei, L., Li, C., Zhao, L., Shen, X., Wang, Y. & Zhang, L. 2014. Asticcacaulis endophyticus sp. nov., a prosthecate bacterium isolated from the root of Geum aleppicum. Int J Syst Evol Microbiol, 64, 3964–3969.

